# A gut-brain axis for aversive interoception drives innate and anticipatory emesis in *Drosophila*

**DOI:** 10.1101/2025.10.01.679515

**Authors:** Rashmi Karunakaran, Prerana Choudhary, Neena Baburaj, Meghna Sawant, Zeyu Chang, Meet Zandawala, Gaurav Das

**Affiliations:** Brain and Feeding Behaviour Laboratory, National Centre for Cell Science, S.P. Pune University Campus, Ganeshkhind, Pune 411007, India; Savitribai Phule Pune University, Ganeshkhind, Pune 411007, India; Regional Centre for Biotechnology, Faridabad; University of Nevada, Reno, Reno, NV, USA; University of Würzburg, Würzburg, Germany

**Keywords:** Vomiting, emesis, Gut-Brain Axis, Memory, Mushroom body, Serotonin, Dopamine, Enteroendocrine cells, Neuropeptides, TrpA1, *Drosophila*

## Abstract

Signals from the gut are increasingly recognized as modulators of brain function and behavior. However, the pathways through which the gut conveys adverse or unpleasant information to the brain are still not well understood. In this study, we identify an aversive gut-brain axis in *Drosophila melanogaster* that detects toxin-induced gut damage and triggers both innate and learned anticipatory emesis (vomiting). After toxin ingestion, reactive oxygen species are produced by midgut enterocytes and detected by the transient receptor potential channel TrpA1 on nearby enteroendocrine cells. This sensing stimulates the release of neuropeptides from enteroendocrine cells, likely representing the gastric malaise flies experience after eating. We show that these neuropeptides act on specific serotonergic and dopaminergic neurons in the brain. These neurons interact with each other and signal to the downstream memory-related mushroom bodies to promote emesis. This circuit not only drives an immediate emetic response but also represents a malaise-driven aversive signal. The signal manifests as the persistent activity of dopaminergic neurons, which reinforces aversive valence to odor cues in the mushroom bodies. Thus, the flies learn that a specific odor predicts the presence of a toxin in food and exhibit anticipatory emesis upon re-exposure to the same odor. Taken together, we have identified an interoceptive signaling pathway that may be conserved for detecting harmful gut conditions and for remembering how to avoid them. Our work offers a mechanistic framework for studying aversive gut-brain communication involved in feeding, metabolism, depression, brain injury, and neurodegenerative diseases.

## Introduction

Incidents in the gut are communicated to the brain, which responds by adjusting gut function accordingly. This complex two-way communication, involving neuronal and hormonal components, is known as the gut-brain axis (Clarke et al., 2024; Mayer et al., 2022). The gut-brain axis governs cognitive, neurological, physiological, and homeostatic functions (Carabotti et al., 2015). Therefore, disruptions in gut-brain communication, including those caused by dysbiosis, have been linked to a variety of conditions such as memory problems, depression, anxiety, neurodegeneration, disorders of gut-brain interactions like irritable bowel syndrome, and metabolic issues (Benakis et al., 2020; Clemmensen et al., 2017; Loh et al., 2024; Mayer et al., 2022; Raber & Sharpton, 2023). Given their clinical and biological significance, interoceptive gut-brain pathways have attracted considerable interest (Alhadeff & Yapici, 2024; W. G. Chen et al., 2021; Craig, 2002). The perception of nutrients after ingestion—such as sugars, amino acids, and salt—through parallel gut-brain axes has been shown to influence diverse behaviors ranging from feeding to sleep in mice and fruit flies (B. Kim et al., 2021; McDougle et al., 2024; Singh et al., 2024; Tan et al., 2020; Titos et al., 2023). In contrast, the gut-brain mechanisms that relay information about post-ingestive gut damage caused by toxins or pathogens remain incompletely understood (Kobler et al., 2020; Xie et al., 2022).

Emesis, or vomiting, is a highly conserved protective response to consuming toxic foods. Besides poisoning, emesis can also occur as a symptom of various conditions, such as microbial infections, cancers, gastrointestinal issues, and from many drugs, including cytotoxic chemotherapeutic agents and anesthetics. Factors like motion sickness, pregnancy, severe emotional stress, and brain injuries also trigger emesis. Interestingly, emesis can also appear as an anticipatory response when re-exposed to environmental cues, such as an odor, that have been linked to previous emetic episodes. Indeed, some cancer patients receiving multiple rounds of chemotherapy develop a learned anticipatory nausea and vomiting (ANV) in response to the sensory context of the drug administration (e.g. smell and sight of the clinic) which can be difficult to treat. ANV is thought to be a case of Pavlovian conditioning, and there are no genetic animal models available (Gupta et al., 2021; Kamen et al., 2014; Zhong et al., 2021).

Despite its biological and clinical importance, the mechanisms of emesis are understood primarily at the level of gross gut-brain anatomy and neurochemistry (Xu et al., 2024; Zhong et al., 2021). This is mainly because widely used vertebrate genetic models, rats and mice, cannot exhibit emesis, likely due to the distinct anatomical shortcomings of their diaphragms and guts. Thus, our current understanding of emesis has been primarily derived from studies of mammals, including dogs, cats, ferrets, and shrews (Horn et al., 2013).

In mammals, the dorsal vagal complex (DVC) of the brainstem serves as a central integration center for aversive visceral signals. The DVC comprises the nucleus tractus solitarius (NTS), the dorsal motor nucleus of the vagus (DMV), and the area postrema (AP). In the gut, enterochromaffin cells detect ingested toxins and release neuroactive substances such as serotonin (5-HT) and substance P (SP) (Bellono et al., 2017; Zhong et al., 2021). These enterochromaffin cells form synapse-like connections with afferent vagal nerve terminals, which express specific receptors including 5-HT3R, 5-HT4R, 5-HT1BR, and NK1 (Heckroth et al., 2021). The signals from the gut travel via the enteric nervous system and visceral afferents—primarily the vagus and splanchnic nerves—which project to the NTS (Spencer & Hu, 2020).

In parallel, the AP, located outside the blood-brain barrier, can directly sense blood-borne toxins, metabolites, and drugs from the circulation and cerebrospinal fluid. These signals also converge onto the NTS. Once activated, the NTS relays information to other brainstem nuclei, including the nucleus ambiguus and the DMV, to coordinate the motor outputs associated with nausea and emesis. This information is presumed to reach higher brain centers for sensory integration and memory formation (Zhong et al., 2021).

Although existing studies have outlined a basic neurophysiological framework, our understanding of emesis, both naive and anticipatory, remains highly fragmented. Fundamental questions persist about how the gut detects harmful agents, the complete repertoire of gut-derived neurohormones involved, and how emetic signals are integrated and processed by the brain. Even less is known about anticipatory emesis, including how aversive memories are formed, reinforced, and retrieved in the absence of an immediate toxin. Together, these gaps highlight the need for a more comprehensive and integrated understanding of the gut-brain axis in emesis.

Here, we use the genetically tractable model organism *Drosophila melanogaster* to define a gut-brain circuit that mediates emetic behavior. We establish that flies show acute emesis following ingestion of toxic substances. Ingested toxins trigger the production of reactive oxygen species (ROS) in midgut enterocytes (ECs), which are detected by the transient receptor potential cation channel A1 (TrpA1) expressed on adjacent enteroendocrine cells (EECs). In response, EECs release multiple neuropeptides that remotely engage discrete populations of serotonergic neurons (5-HTNs) and dopaminergic neurons (DANs) in the fly central brain. 5-HTNs and DANs reciprocally regulate each other to modulate emesis, and in turn, signal to the mushroom bodies (MBs), centers of associative learning and memory in the fly brain, driving naive emesis. Interestingly, the same pathway mediates aversive learning, enabling flies to show anticipatory emesis when re-exposed to an odor previously paired with toxin ingestion and gut distress. Our work establishes the fruit fly as a genetic model for studying aversive, interoceptive gut-to-brain pathways, especially in the context of naive and learned emesis.

## Results

### Ingestion of toxins induces emesis in flies

Toxin ingestion is known to trigger emesis in many animals (Wickham, 2020; Xu et al., 2024; Zhong et al., 2021). We first aimed to determine whether the ingestion of toxic and/or bitter compounds, such as copper sulfate, nicotine salts, denatonium, caffeine, coumarin, and lithium chloride (Weiss et al., 2011), also leads to emesis in flies. To test this, we fed 24-hour starved, 4-5 days old, mixed-sex flies (∼5 flies per group) with a mixture of 750 mM sucrose and toxins in custom-designed arenas (Fig. 1A). Sucrose was added to mask immediate taste-driven rejection of toxic compounds and to mimic unintentional ingestion of toxins with food. We added a blue dye to the mix to visually observe feeding and emesis. Indeed, toxin ingestion resulted in the oral deposition of blue droplets within the arenas (Fig. 1A, Fig. S1-1). Assay time was set to 30 minutes, as we found that flies consuming sucrose-toxin mixtures, except for denatonium, rarely defecated during this period (Supplementary Video 1; Fig. S1-2A) Emetic droplets were either counted manually or analyzed from recorded images and videos. We measured the extent of emesis as the number of emetic droplets per fly (droplets/fly) from each arena (Fig. 1B).

**Figure 1.**
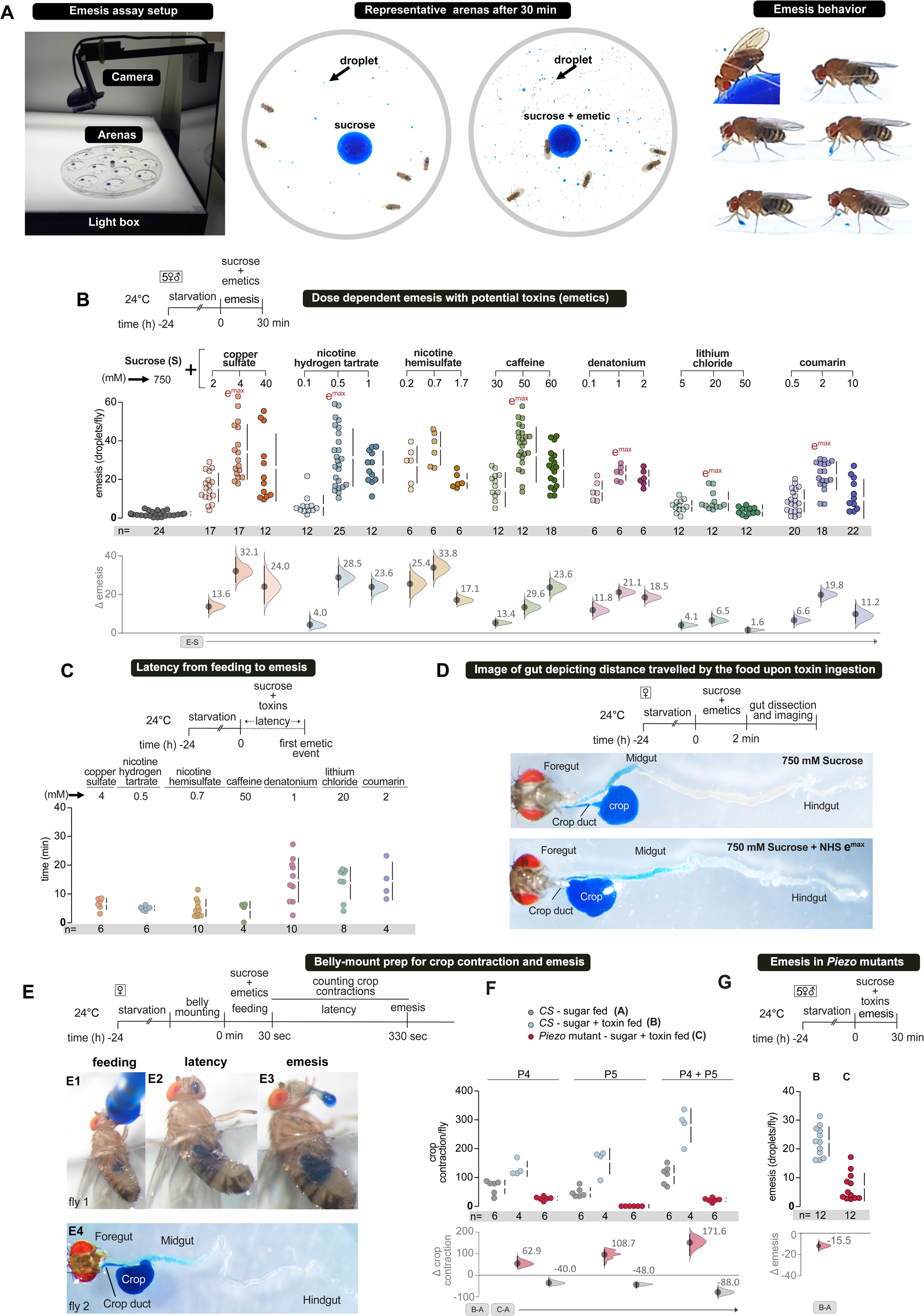
*Drosophila* show dose-dependent, delayed, and post-ingestively triggered emesis following toxin ingestion. **(A)** Setup for video-recording the emesis assay (*left*), representative images of control (sucrose only) and experimental (sucrose + emetic) arena showing emetic droplets at the end of a 30 min assay period (*middle*), and close-up of a fly from feeding to vomiting (*right*). **(B)** Mixed sex flies exhibit dose-dependent emesis in response to multiple potential emetics. Flies were fed a mixture of 750 mM sucrose + emetics + blue dye. Emesis is quantified as the number of blue emetic droplets per fly. Emesis increases with toxin concentration up to an optimal dose (e^max^), beyond which it decreases. **(C)** Individual flies typically show a temporal latency of ∼ 5 minutes or more between ingestion of sugar-toxin mixtures and showing emesis, indicating that emesis is likely triggered in the gut after ingestion. **(D)** Distance travelled by the food after 2 minutes of feeding. The ingested food, sucrose only (above) and sucrose+ toxin mixture (below), reaches the midgut of flies within this time **(E)** Measuring food passage into the gut and crop contractions in immobilised flies using the bellymount preparation. All stages of feeding, latency, and emesis can be observed in bellymounted flies (E1-E3). Toxic food is clearly visible in the midgut a minute after feeding (E4). **(F)** Quantification of P4 and P5 muscle contractions from bellymounted *CS* and *piezo* mutant flies over a period of 5 minutes after feeding. **(G)** Emesis is drastically reduced in *piezo* mutant flies.

We observed that flies generally showed a dose-dependent increase in emesis with rising toxin concentrations in the mixture, up to a certain point (Fig. 1B; note the e^max^). However, at higher relative toxin levels, emesis tended to decrease (Fig. 1B). Next, we confirmed the masking of the aversive tastes of nicotine salts and copper sulfate with the sucrose carrier using the proboscis extension response (PER). At the e^max^ concentration, these compounds were unable to suppress PER with 750 mM sucrose alone (Fig. S1-2B), indicating that the flies probably did not sense the toxin in the mix. Adding toxins at concentrations higher than e^max^ began to suppress sucrose PER, suggesting that the flies were now detecting the toxins (Fig. S1-2B). These higher concentrations of nicotine salts and copper sulfate also reduced feeding in the blue dye assay (Fig. S1-2C), likely contributing to the dip in emesis observed (Fig. 1B). Moving forward, we primarily used nicotine hydrogen tartrate (NHT), nicotine hydrogen sulfate (NHS), and copper sulfate in all experiments.

Although flies did not exhibit aversive PER responses to mixtures of sucrose and toxins at the periphery, we could not rule out the possibility that bitter gustatory receptor neurons (GrNs) physiologically respond to toxins to induce emesis. To explore this, we tested for emesis after feeding flies only 750 mM sucrose. At the same time, we activated bitter GrNs labeled with *Gr66a-GAL4* by driving the expression of the *UAS-dTrpA1* cation channel (Hamada et al., 2008; Kang et al., 2010). The experimental flies did not show increased emesis compared to their parental controls (Fig. S1-2D).

Blocking bitter GrNs transiently, using the *UAS-Shibire^ts1^* (Kitamoto, 2001) transgene in flies fed only sugar or sugar-toxin mixtures at e^max^, did not affect emesis either (S1-2E). Hence, bitter GrNs do not appear to be involved in inducing emesis.

### Toxins reach the midgut before intense crop contractions and emesis

We reasoned that emesis should occur without much delay after contact with food if triggered by peripheral bitter taste. On the other hand, a significant latency between food ingestion and emesis would indicate that the emesis was initiated well after ingestion. To distinguish between these possibilities, we measured the time between ingestion and emesis in flies by analyzing video recordings of their behavior. We found a significant delay between ingestion and emesis for all compounds tested. Mean latency times ranged from approximately 5 to 15 minutes (Fig. 1C). This extended delay strongly suggested that emesis was triggered well after the food had been ingested and entered the gut.

To identify the part of the gut where toxins are detected, we examined how toxin-containing food is transported into the gut before being expelled through vomiting. We fed individual flies a blue-dyed sucrose + NHS solution for 2 minutes inside a small tube, then immediately dissected the gut. In all samples, we observed the presence of blue dye in the midgut. The movement of blue dye was similar in control guts fed only sucrose (Fig. 1D; representative images). This shows that toxic food reached the midgut well before the typical latency for NHS-induced vomiting (Fig. 1C).

The insect crop has been linked to regurgitation (Stoffolano & Haselton, 2013). To study the role of crop contractions in vomiting, we used the belly-mount preparation (Koyama et al., 2020). Single flies were fed a blue-dyed emetic mixture in three short bouts lasting 20-30 seconds under a stereo microscope. We counted the contractions of the P4 and P5 crop muscles for an additional 5 minutes (Fig. 1E; schema). We observed food moving into the crop almost immediately (Fig. 1E1). The P4 muscle started contracting around 83 ± 4 seconds (mean ± s.d.). These initial P4 contractions facilitate the passage of food into the midgut and are observed in both experimental and control flies. P4 contractions increase significantly in experimental flies over the next few minutes (Fig. 1F). The P5 muscle initiates contractions around 106 ± 16.4 seconds (mean ± s.d.) after feeding begins in the experimental group and is much more frequent and intense than in control flies (Fig. 1F, Supplementary video 2). In belly-mounted flies, droplet expulsion or vigorous proboscis extensions were observed after a latency period (Fig. 1E2-1E3, Supplementary video 2). These intense proboscis extensions appeared in experimental flies approximately 295 ± 111 seconds after feeding started. Based on this data and the direct observation of belly-mounted flies, we concluded that crop contractions preceded and facilitated emesis.

Next, to determine whether food passes to the midgut before intense crop contractions in belly-mounted flies, we dissected guts 90 seconds after feeding started, coinciding with the initial P4 contractions. We clearly observed blue dye in the midgut of all the dissected crops, suggesting that food passed to the midgut well before intense crop contractions and emesis (Fig. 1E4; representative image).

Overall, the contractions of P4 and P5 were nearly three times higher in toxin-fed flies compared to control sugar-fed flies (Fig. 1F). In *Piezo* mutant flies (Kim et al., 2012; Wang et al., 2020), crop contractions after feeding the NHS were almost absent (Fig. 1F). As a result, these flies could not vomit, further emphasizing the importance of proper crop function in emesis (Fig. 1G). Since a non-permeable cuticle lines the foregut (Miguel-Aliaga et al., 2018), we hypothesized that toxins would likely be detected in the midgut.

### Toxin ingestion triggers ROS production from ECs, which is detected by the dTrpA1 channel on EECs

The fly gut is an epithelial tube surrounded by muscles and connected by nerves and trachea. It is divided into three principal regions: the foregut, midgut, and hindgut. In adult flies, the midgut is further divided into five distinguishable regions (R1 to R5), as shown in Fig. 2A. Four primary cell types are found in the midgut: regenerative intestinal stem cells (ISCs), transit-amplifying enteroblasts (EB), nutrient-absorbing ECs, and sparse, secretory EECs (Fig. 2A) (Buchon et al., 2013; Miguel-Aliaga et al., 2018).

**Figure 2.**
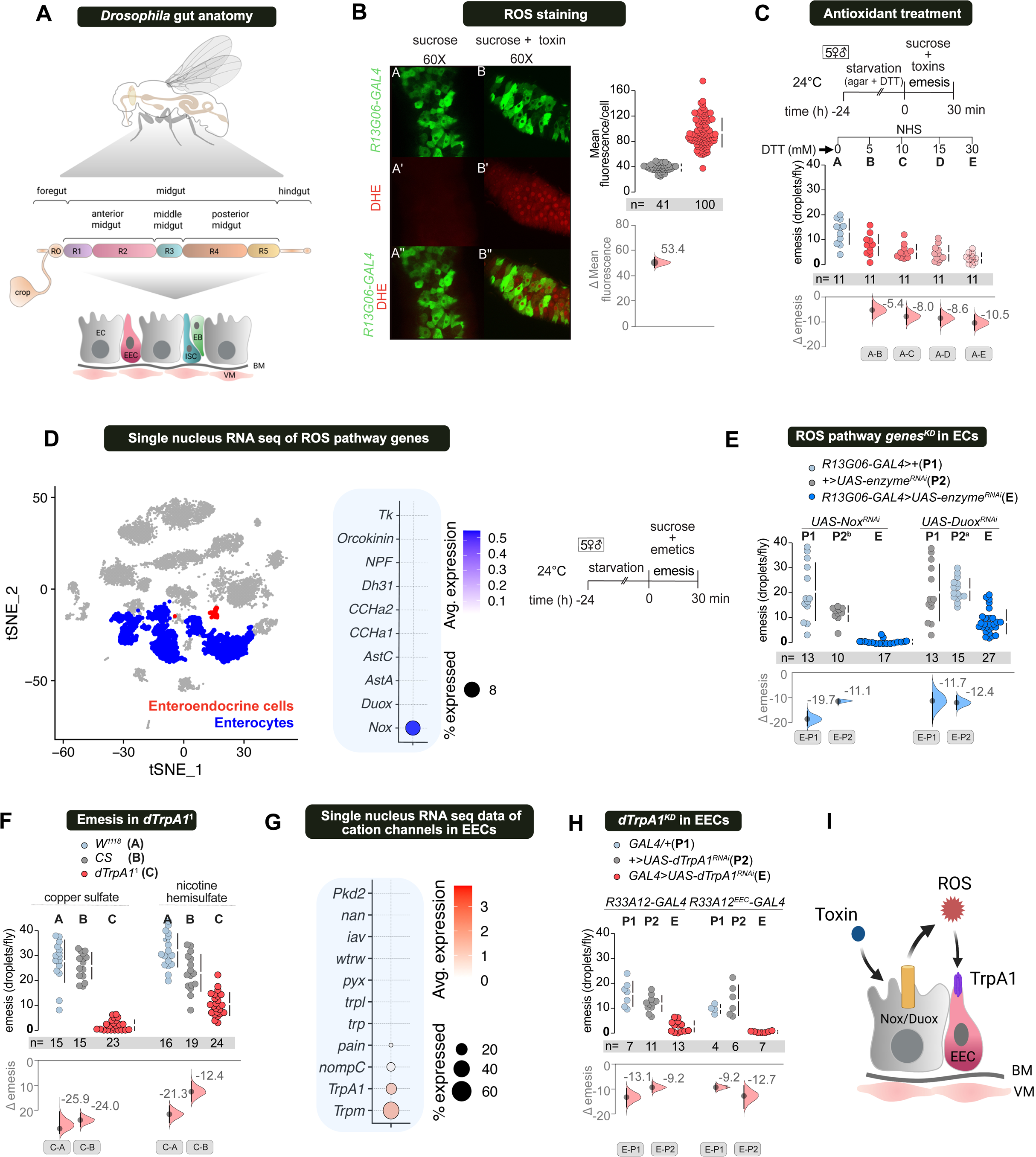
ROS generated by ECs after ingestion of emetics is detected by dTrpA1 channel on EECs. **(A)** Cellular organization of the *Drosophila* midgut, showing interstitial stem cells (ISCs) and differentiated cells. Note the epithelial enterocytes (ECs), the most common cell type, and the midgut-specific secretory enteroendocrine cells (EECs). **(B)** DHE staining reveals that guts fed with emetics have significantly higher levels of reactive oxygen species (ROS) production compared to control guts fed only sucrose. **(C)** Loading flies with the antioxidant dithiothreitol (DTT) while they are starved before the emesis assay dose-dependently reduces emesis. **(D)** Midgut ECs and EECs separated based on single nucleus expression profiling from (H. Li et al., 2022) (left panel). Quantitative single nucleus RNA sequencing data reveal the presence of neuropeptides and ROS pathway genes in the ECs (right panel). Neuropeptide expressions are below detection levels in the ECs. While the ROS-processing enzyme NADPH oxidase (*Nox*) is abundant, the expression of Dual Oxidase (*Duox*) is below the detection limit. **(E)** However, UAS-RNAi knockdown of both *Nox* and *Duox* in an EC-specific line *R13G06-GAL4* strongly inhibits emesis.**(F)** *dTrpA1* mutant flies (*dTrpA1^1^*) show severely reduced emesis with both copper sulfate and nicotine hemisulfate. **(G)** Quantitative single nucleus RNA sequencing data for cation channel genes in the ECs (H. Li et al., 2022). **(H)** RNAi-mediated knockdown of *dTrpA1* in EECs, driven by *R33A12-GAL4* and its EEC-specific variant *R33A12-GAL4^EEC^*, also significantly decreases emesis. **(I)** Summary figure: Toxin triggers EC-mediated ROS generation, which dTrpA1 senses in EECs to induce emesis.

Besides aiding digestion and nutrient absorption, the gastrointestinal tract plays a vital role in recognizing pathogens and toxins (Bonnay et al., 2013; Hayakawa et al., 2004; Iwasaki & Medzhitov, 2010; Kuraishi et al., 2011; Lemaitre & Hoffmann, 2007; O’Neill et al., 2013; Tellam, 1996).

*Gr* promoter-driven GAL4 lines are expressed in specific subsets of midgut EECs, suggesting that the corresponding *Grs* may also be present there (Park & Kwon, 2011). Gr66a or Gr33a are involved in detecting various bitter compounds, including copper sulfate and nicotine (Rimal & Lee, 2019; Xiao et al., 2022). To explore the potential role of bitter GrNs in the gut for inducing emesis, we knocked them down using UAS-RNAi constructs in a relatively broad group of EECs labeled with the *R33A12-GAL4* line (Fig. S2-1A) (Lim et al., 2021). While knocking down *Gr66a* did not appear to cause any defect in emesis compared to controls, knocking down *Gr33a* seemed to cause a slight decrease in emesis with NHS but not copper sulfate (Fig. S2-1A). Overall, our data suggest that gut-expressed Gr66a and Gr33a are unlikely to be involved in sensing copper sulfate and nicotine to trigger emesis.

Previous studies in flies suggest that enteric bacterial infection or exposure to potentially toxic chemicals can trigger the release of ROS from ECs into the gut lumen (Iatsenko et al., 2018; Jones et al., 2013; Kuraishi et al., 2013; Singh et al., 2009; Yao et al., 2016).

Consequently, we questioned whether toxin-induced vomiting is mediated by ROS production and detection. To address this, we measured ROS levels in the fly guts after toxin ingestion using a ROS-specific DHE dye stain. Flies that were allowed to feed on a sugar-toxin mixture for 10 minutes showed higher ROS levels in their guts compared to control flies fed only sucrose (Fig. 2B). Preloading the flies with the ROS-neutralizing antioxidant dithiothreitol (DTT) for a day before the assay reduced vomiting in a dose-dependent manner, further implicating ROS (Fig. 2C). Feeding assay performed in these flies showed comparable feeding levels as that of its control group, suggesting that the reduced vomiting was not a result of reduced feeding (S2-1D).

To identify genes expressed in ECs that could be involved in toxin detection and ROS production, we mined single-nucleus transcriptomes of midgut cells for genes related to ROS generation (H. Li et al., 2022)(FlyCellAtlas). Our analysis revealed that the ROS-producing gene NADPH oxidase (*Nox*) is expressed in *Drosophila* ECs (Fig. 2D). To determine if Nox expression in ECs is necessary for emesis, we knocked it down in ECs labelled by *R13G06-GAL4* (Fig. S2-2A) (Lim et al., 2021) and *NP1-GAL4*. This resulted in a drastic reduction of emesis (Fig. 2E, S2-1B). Not detecting other ROS-producing genes in the EC single-nucleus transcriptome data does not mean that they are not expressed. Hence, we also used EC-specific RNAi-mediated knockdown of the other essential ROS-producing gene, *Duox* (Jones et al., 2012; Kim & Lee, 2014; Kuraishi et al., 2013). This manipulation also strongly reduced emesis (Fig. 2E, S2-1B), confirming that ROS production by ECs triggers emesis.

Recent reports indicate that TrpA1 functions as a receptor for ROS, chemical irritants, and pathogens (Du et al., 2016; Kang et al., 2010; Kim et al., 2010; Y. Li et al., 2023). Therefore, we first tested for emesis in *dTrpA1* mutant flies (*dTrpA1^1^*). We found that these flies exhibited a significant reduction in emesis compared to control wild-type flies (Fig. 2F). Single-nucleus transcriptome analysis of genes in the TRP superfamily revealed that *TrpA1* is highly expressed in midgut EECs (Fig. 2G) (Gong et al., 2023; H. Li et al., 2022). Hence, we used RNAi to knockdown *dTrpA1* in EECs. Knockdown of the *dTrpA1* gene in EECs using *R33A12-GAL4* and *R33A12-GAL4^EEC^*, an EEC-specific driver combined with *nSyb-GAL80* (Fig. S2-2B), resulted in nearly complete suppression of emesis (Fig. 2H). We also knocked down *dTrpA1* in ECs using *R13G06-GAL4* and *NP1-GAL4* driver lines. We observed no reduction in emesis (S2-1C). Independently, we confirmed that the reduction in emesis was not due to decreased feeding in *dTrpA1^1^ mutant* flies or *dTrpA1* knockdown flies (Fig. S2-1E, F). On the contrary, *dTrpA1* knockdown in EECs results in increased feeding (Fig. S2-1E, F). These findings strongly suggest that EC-generated ROS are sensed by dTrpA1 in EECs to mediate emesis (Fig. 2I).

### Neuropeptides AstA, AstC, Tk, DH31, and CCHa2 from EECs mediate emesis

In mammals, activating TrpA1 with electrophiles causes the release of 5-HT from enterochromaffin cells. This 5-HT release represents the aversive signal from the gut (Bellono et al., 2017; Nozawa et al., 2009). However, analysis of single-nucleus gene expression data for the midgut EECs does not show any detectable expression of tryptophan hydroxylase neuronal (*Trhn*), which is involved in 5-HT biosynthesis, and vesicular monoamine transporter (*Vmat*), which is engaged in packaging monoaminergic neurotransmitters like 5-HT into vesicles for later release (data not shown). Additionally, knocking down *Trh* in fly EECs using *R33A12-GAL4* does not affect emesis (Fig. S3-1A). Together, these data suggest that unlike mammals, 5-HT is not produced in *Drosophila* EECs, and another signal from the EECs likely mediates emesis.

The activation of *dTrpA1* triggers neuropeptide release in flies (Benguettat et al., 2018; Ogawa et al., 2016). Fly midgut EECs produce neuropeptides or neurohormones, with their expression varying across the midgut zones R1 to R5 (Fig. 3A) (H. Li et al., 2022) (Nässel & Zandawala, 2019; Reiher et al., 2011). To determine whether neuropeptide processing in EECs is essential for emesis, we used RNAi to knockdown the prohormone convertase *amon* using the EEC driver *R33A12-GAL4*. Amon cleaves peptide hormone precursors to produce active forms, and the absence of amon in EECs significantly decreases peptide production (Reiher et al., 2011; Wegener & Veenstra, 2015). We found that knocking down *amon* in EECs considerably lowered emesis, indicating that peptide processing, and thus neuropeptide production, in EECs is necessary for this behavior (Fig. S3-1B).

**Figure 3.**
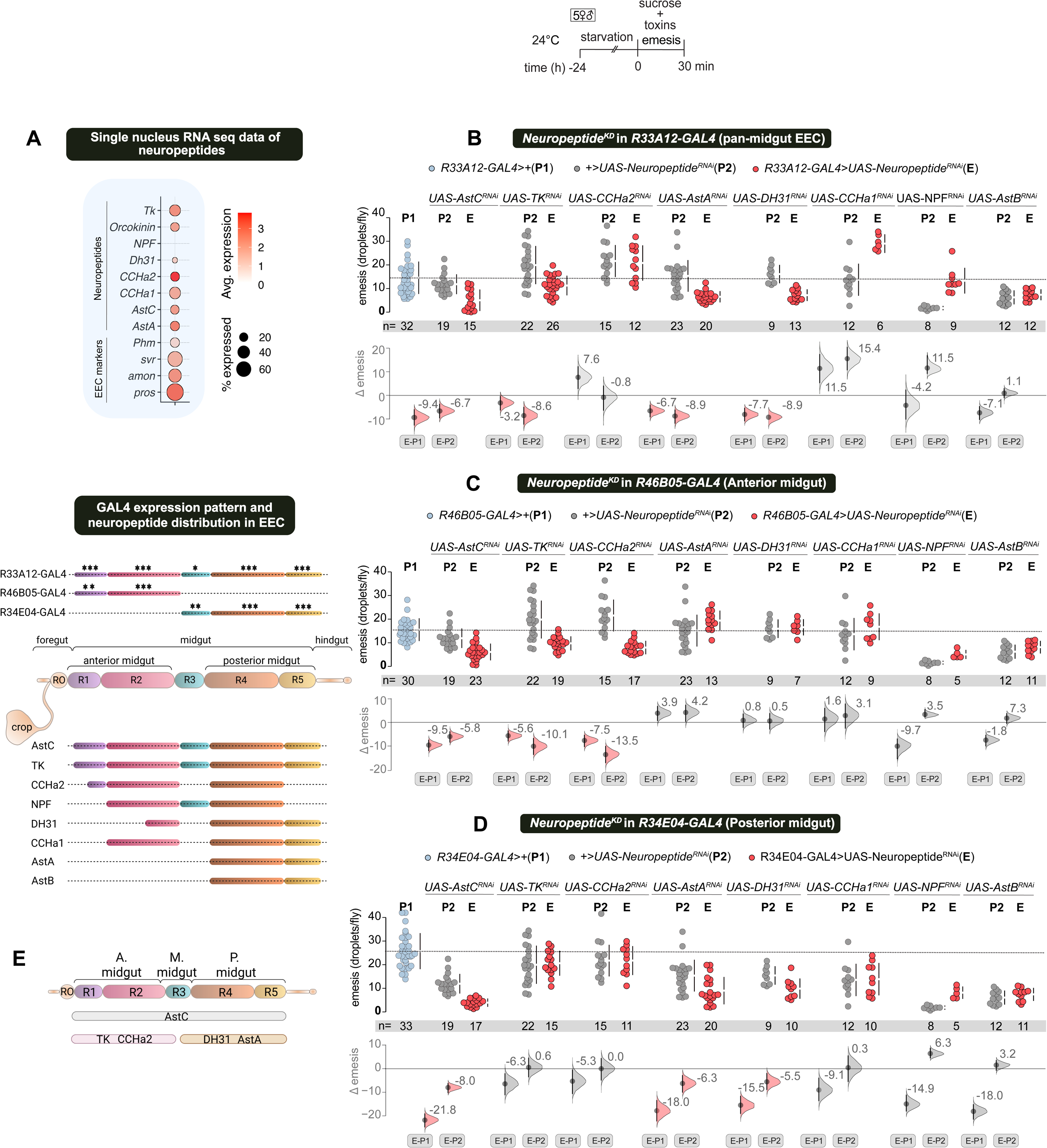
AstC, from the whole midgut, anteriorly expressed Tk and CCHa2, and posteriorly expressed AstA and Dh31, facilitates emesis. **(A)** Midgut ECs and EECs are distinguished based on single nucleus RNA expression profiling from (H. Li et al., 2022) (top panel). Quantitative single-cell RNA sequencing data reveal the expression of neuropeptide and neuropeptide processing genes in the EECs (middle panel). Zone-specific neuropeptide expression is observed in the fly midgut, along with the coverage of GAL4 lines used to inhibit neuropeptide expression via RNAi (bottom panel). **(B)** Inhibiting candidate neuropeptide expression in *R33A12-GAL4*, the broadest midgut GAL4 line used, suggests that AstC, AstA, Tk, and Dh31 may play roles in mediating emesis. **(C)** Inhibition in *R46B05-GAL4*, an anterior-biased midgut GAL4 line, indicates that anteriorly expressed AstC, Tk, and CCHa2 could be involved in emesis. Knockdown of *AstA* and *Dh31*, which are not expressed anteriorly, does not reduce emesis with this line. **(D)** Inhibition in *R34E04-GAL4*, a posterior-biased midgut GAL4 line, points to a role for posteriorly expressed AstA and Dh31 in emesis. Tk produced from the posterior midgut does not appear relevant for emesis. Finally, *AstC* inhibition decreases emesis with this line. **(E)** Overview of neuropeptide expression from the midgut that mediates emesis. The dashed line in the graphs indicates the mean of the common control group P1.

Next, to identify the specific neuropeptides involved, we used RNAi-mediated knockdown of all candidate peptides in three different GAL4 lines: *R33A12-GAL4* broadly labels R1-R5 EECs; *R46B05-GAL4* is biased toward the anterior midgut and primarily labels R1-R2 EECs; and *R34E04-GAL4* is biased toward the posterior midgut and labels R4-R5 EECs (Fig. 3A and Fig. S2-2B, C, D). We found that knockdown of *allatostatin C* (*AstC*) with all three driver lines reduced emesis in flies, indicating its widespread role along the midgut (Fig. 3B-D). While tachykinin (Tk) and CCHamide 2 (CCHa2) peptides are expressed across all midgut zones, their knockdown in the anterior-biased line *R46B05-GAL4*, but not in the posterior-biased *R34E04-GAL4*, affected emesis (Fig. 3C, D). Allatostatin A (AstA) and diuretic hormone 31 (Dh31) are known to be expressed in the posterior midgut, and knocking them down in the posterior zone, but not the anterior zone, reduced emesis (Fig. 3C, D). To further attribute the role of these neuropeptides solely in EECs, we used the same three EEC drivers (combined with *nSyb-GAL80* to suppress neuronal expression) for RNAi knockdown. These manipulations reduced emesis in all cases, confirming that EEC-derived neuropeptides regulate emesis (Fig. S3-1C, D, E).

Gut-derived neuropeptides, including AstA, AstC, Tk, CCHa2, and Dh31, are also known to influence feeding and metabolic homeostasis (Wu et al., 2020; Zhou et al., 2020). In the emesis assay, the flies are hungry, and the majority of their feeding on sugar-emetic mixtures occurs in their initial feeding bout. Therefore, we conducted a 3-minute blue dye feeding test to confirm that the reduced vomiting observed in previous experiments was not due to decreased feeding during the assay’s feeding phase, which could have been caused by neuropeptide knockdown. However, we saw no reduction in feeding on a timescale relevant to our assay (Fig. S3-1F, G, H). Overall, these results suggest that toxin-triggered activation of *dTrpA1* in EECs induces vomiting via multiple neuropeptides.

### Serotonergic and dopaminergic neurons are required for emesis

Both 5-HTNs and DANs are involved in mammalian emesis (Belkacemi & Darmani, 2020; Johnston et al., 2014; Minami et al., 1996). 5-HTNs have been proposed to be activated by aversive events (Deakin & Graeff, 1991) and shown to influence aversive learning (Chakraborty et al., 2019; Sitaraman et al., 2017; Tortora et al., 2023; M.-S. Wu et al., 2023). 5-HT levels increase in the mammalian brain after various stressors (Adell et al., 1997; Yoshioka et al., 1995). In insects, such as honeybees, 5-HT seems to represent post-ingestive malaise (Wright et al., 2010). Hence, while our earlier experiments ruled out the involvement of peripheral 5-HT signaling from the gut in inducing emesis, central brain 5-HT neuronal signaling could still be involved. DANs have been clearly implicated in the representation of both aversive punishment and appetitive reward in both mammals and insects (Adel & Griffith, 2021; Schultz, 2010; Waddell, 2013).

Therefore, to investigate the role of 5-HT and dopamine (DA) in fly vomiting, we conducted a small-scale pharmacological screen using known inhibitors of 5-HT and DA receptors (5-HTRs and DARs). We observed that the administration of 5-HTR and DAR antagonists indeed affected emesis in wild-type flies (Fig. S4-1A), indicating that neuronal 5-HT and DA signaling might be involved

To test this prediction, we conducted a targeted neurogenetic screen by acutely silencing specific groups of 5-HTNs and DANs using the *UAS-shibire^ts1^* transgene (Fig. 4A). Among the different subsets of 5-HTNs tested, acute silencing of 5-HTNs marked by *TRH::T2A-GAL4* significantly impaired emesis in flies (Fig. 4A, Fig. S4-1B) .

**Figure 4.**
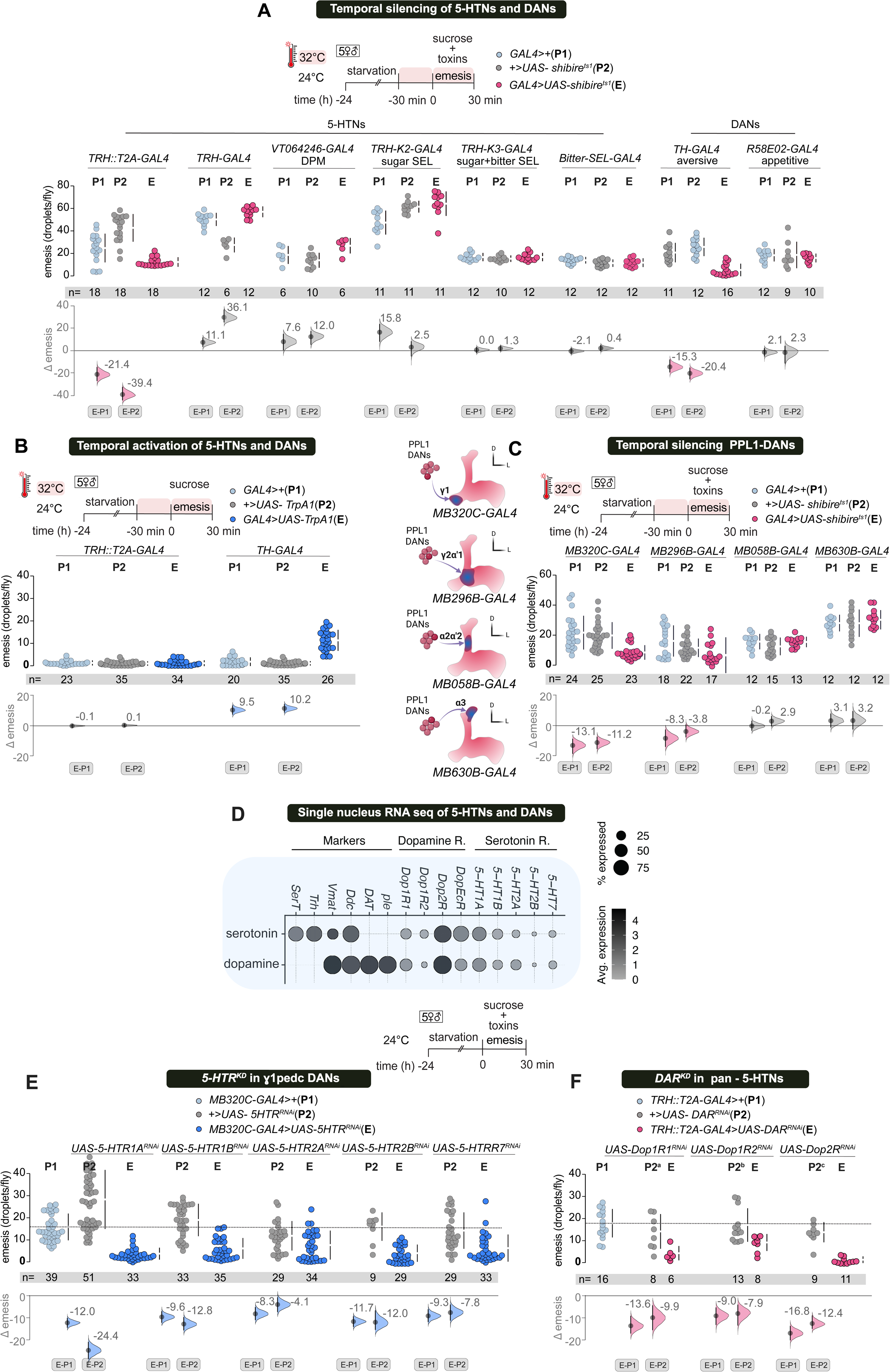
Mushroom Body innervating Dopaminergic neurons (DANs) receive input from Serotonergic neurons (5-HTNs) and signal aversive tone for emesis. **(A)** Transient thermogenetic silencing of various subpopulations of 5-HTNs and DANs during emesis assay. Disruption of emesis was observed upon silencing *TRH::T2A-GAL4* (5-HTNs) and *TH-GAL4* (DANs) labeled neurons. **(B)** Artificial thermogenetic activation of *TH-GAL4* labeled DANs induces emesis in flies fed only sucrose, indicating that they signal an aversive valence related to emesis. In contrast, activating *TRH::T2A-GAL4* 5-HTNs did not increase emesis in flies fed only sucrose.**(C)** Behavioral-genetic screen to identify the PPL1 DAN subset necessary for emesis reveals that transient silencing of the mushroom body heel innervating PPL1-γ1pedc DANs (*MB320C-GAL4*) using *UAS-Shibire^ts1^*drastically reduced emesis. Silencing PPL1-γ2ɑ’1 DANs (*MB296B-GAL4*) caused a moderate reduction. Conversely, silencing ɑ2ɑ’2 (*MB058B-GAL4*) or ɑ3 (*MB630B-GAL4*) DANs had no effect. **(D)** Single nucleus transcriptomics data from (H. Li et al., 2022) suggest that 5-HTNs and DANs express nearly all serotonergic and dopaminergic receptors. **(E)** RNAi knockdown of *5-HTR1A*, *5-HTR1B*, *5-HTR2B*, and *5-HTR7* in PPL1-γ1pedc DANs (*MB320C-GAL4*) all reduced emesis to varying degrees. **(F)** RNAi knockdown of dopamine receptors (DARs) *Dop1R1*, *Dop1R2*, and *Dop2R* in 5-HTNs labeled by the broad *TRH::T2A-GAL4* also reduced emesis levels to different extents.

Additionally, chronic inhibition of *TRH::T2A-GAL4* labeled neural activity, using *UAS-Kir2.1*, an inward-rectifying potassium channel transgene (Baines et al., 2001), also markedly decreased emesis (Fig. S4-1C). The *TRH::T2A-GAL4* is broad, and our analysis with anti-5-HT co-staining shows that it labels nearly all 5-HTNs and also some neurons not labeled by 5-HT (*data not shown*). Neurons in this *GAL4* line also have mostly axonal projections on the foregut-midgut junction, the crop, proventriculus/cardia, and the anterior midgut (Fig. S4-1B). Silencing the dorsal paired medial (DPM) neurons, which are part of the *TRH::T2A-GAL4* expression pattern, does not suppress emesis (Fig. 4A, Fig. S4-1B). Nutrient sugar and bitter compounds activate specific 5-HT neurons (sugar and bitter-SELs) in the fly brain. The activity of bitter-SELs triggers crop contraction (Yao & Scott, 2022). While the bitter-SELs are included in *TRH::T2A-GAL4*, neither acute nor chronic silencing of these neurons affected emesis. (Fig. 4A, Fig. S4-1C). Surprisingly, emesis was also unaffected when another broad TRH-GAL4 (Pooryasin & Fiala, 2015) labeled group of neurons was inhibited. Thus, although these experiments exclude certain subsets of 5-HTNs, the specific 5-HTNs that regulate emesis remain unidentified.

Similarly, silencing tyrosine hydroxylase (TH) promoter driven *TH-GAL4*, which labels a large subset of DANs including those involved in aversive reinforcement to MB neuropils, also significantly reduced emesis. Silencing the dopaminergic protocerebral anterior medial (PAM) neurons, which innervate the MB and are involved in reward signaling, which are labeled by *R58E02-GAL4*, did not affect emesis (Fig. 4A). Chronic inhibition of the *TH-GAL4* DANs, using *UAS-Kir2.1*,also decreased emesis (Fig S4-1C).

Having implicated 5HTNs and DANs, we wanted to confirm that the reduction in emesis was explicitly due to 5-HT and DA release respectively, and not another neurotransmitter coexpressed in these neurons. For this, we performed an RNAi-mediated knockdown of the *Trh* gene and also the serotonin transporter (*SerT*) that reuptake 5-HT in *TRH::T2A-GAL4*. Our manipulations drastically reduced emesis (Fig.S4-2A). To examine the role of DA in emesis, we drove RNAi-mediated knockdown of the DA-specific biosynthetic enzyme *TH*, in *TH-GAL4*, and we observed a decrease in emesis (Fig.S4-2B). We conducted an acute 3-minute blue-dye feeding assay in the relevant fly strains to confirm that the reduction in emesis was not due to reduced feeding (Fig. S4-2D, E, F). These results indicate that similar to mammals 5-HT from 5HTNs and DA from DANs is required for emesis.

We then aimed to determine whether the activity of the identified neuronal groups was sufficient to cause emesis. We found that artificial thermogenetic activation of *TH-GAL4* neurons with *UAS-TrpA1* expression in hungry flies that were fed only sucrose at 32 °C increased emesis. However, activation of *TRH::T2A-GAL4* under identical conditions did not show any emesis. (Fig. 4B). This implied that neurons labeled by *TH-GAL4* were sufficient to drive the whole regurgitative/emetic program downstream, which includes regulating food movement from the crop towards the proboscis, instead of the midgut. However, while neurons labeled by *TRH::T2A-GAL4* were required for emesis, unlike TH-GAL4, their artificial activation was probably not able to drive the emetic program in the gut.

The *TH-GAL4* labels several clusters of DANs. To identify the specific DANs involved in emesis at the cellular level, we acutely inhibited synaptic transmission from subsets of the *TH-GAL4* neurons using *shibire^ts1^*. We utilized previously characterized transgenic lines in which specific subsets of *TH-GAL4* are labeled (Liu et al., 2012). Silencing of *TH-C’-GAL4*, which includes the PPL2, PPM2, and PAL clusters, did not affect emesis. Conversely, silencing *TH-D’-GAL4*, comprising the PPL1, PPM2, and PPM3 neurons, as well as *TH-D4-GAL4*, which labels a more restricted set of PPL1, PPM2, and PPM3 neurons, significantly reduced emesis in flies compared to controls (Fig. S4-2C). The PPL1 neurons, common to both lines, consist of 12 neurons that innervate the MBs and provide aversive reinforcement for olfactory memory formation (Aso et al., 2012).

Therefore, we examined the role of PPL1 subsets in emesis. Acute silencing of neurons with *Shibire^ts1^* using the refined *MB320C-GAL4* pattern, which labels PPL1-γ1pedc, largely eliminated emesis, whereas silencing *MB296B-GAL4*, which labels γ2ɑ’1 DANs, caused a small reduction. In contrast, silencing *MB058B-GAL4*, labeling ɑ2ɑ’2 DANs, and *MB630B-GAL4*, labeling ɑ3 DANs, showed no decrease in emesis (Fig. 4C). We conclude that the DANs innervating the MB-heel (PPL1-γ1pedc) were primarily needed for emesis, with a minor contribution from the DANs innervating the proximal vertical MB lobe (γ2ɑ’1).

### 5-HTNs and DANs exhibit reciprocal regulation of emesis

Our results show that silencing neurotransmission from both 5-HTNs and DANs drastically reduces emesis. Hence, these circuits and/or signaling pathways could interact to regulate emesis. To explore synaptic connectivity between these neurons, we leveraged the FlyWire adult female brain connectome (Dorkenwald et al., 2024; Schlegel et al., 2024). We first identified neurons known to express 5-HT based on experimental evidence (Fig. S4-3A) as well as PPL1 DANs (Fig. S4-3B) which have been annotated in a publicly available dataset (see materials and methods for details). Mapping the input and output synapses for these two sets of neurons (Fig. S4-3C,D) shows that these neurons could interact in the heel of the MB or in the superior medial protocerebrum. To examine synaptic interactions between these neurons, we calculated the influence that starter neurons have on receiving neurons (Bates et al., 2025). We observed that 5-HTNs have a strong influence on PPL1 DANs (Fig. S4-3E), while PPL1 DANs have a moderate influence on 5HTNs. The presence or absence of bitter-SELs in the starter neurons has negligible effect on downstream neuron classes examined here (Fig. S4-3F). The differences in influence can be attributed to the differences in number of starter neurons and/or the strength of synaptic connections. In addition to synaptic connections, these neurons could signal extrasynaptically with 5-HT and DA in a paracrine fashion.

Multiple 5-HTRs and DARs are present in *Drosophila* (Blenau & Thamm, 2011; Hearn et al., 2002; Kasture et al., 2018). Single-nucleus transcriptomic analysis (H. Li et al., 2022) clearly demonstrates that 5-HTNs and DANs express receptors for both 5-HT and DA (Fig. 4D), suggesting that these neurons could communicate with each other via paracrine signaling. To test this hypothesis, we first performed RNAi-mediated knockdown of *5-HTRs* in emesis-relevant DANs. Knockdown of *5-HTR1A*, *5-HTR1B*, *5-HTR2B*, and *5-HTR7* in *TH-D4-GAL4*, which includes PPL1 DANs, resulted in a considerable reduction in emesis (Fig. S4-3G). Similarly, knocking down these receptors in PPL1, γ1pedc using *MB320C-GAL4* also decreased emesis (Fig. 4E). This indicates that 5-HT signaling may act upstream of the aversive DANs and be part of the gut-to-brain afferent signaling pathway. Conversely, knocking down DARs, Dop1R1, Dop1R2, and Dop2R in *TRH::T2A-GAL4* also reduces emesis (Fig. 4F). Overall, these findings strongly suggest that, similar to mammals, emesis in flies requires the function of both 5-HT and DA. Furthermore, in flies, emesis-relevant 5-HTNs and DANs likely form an afferent, interoceptive axis that conveys the aversive valence of toxin ingestion, most likely to the MB circuitry.

### Gut neuropeptides communicate with 5-HTNs and DANs for emesis

Neuropeptides released from the gut EECs can signal to central brain neurons following their release into the hemolymph (J. Chen et al., 2024; Lin et al., 2022). We wondered whether gut-released neuropeptides involved in emesis, namely AstA, AstC, CCHa2, DH31, and Tk, signal to the 5-HTNs and DANs via the hemolymph. Indeed, single nucleus RNA sequencing analysis of 5-HTNs and DANs showed that they express receptors for the neuropeptides mentioned above (H. Li et al., 2022) (Fig. 5A).

**Figure 5.**
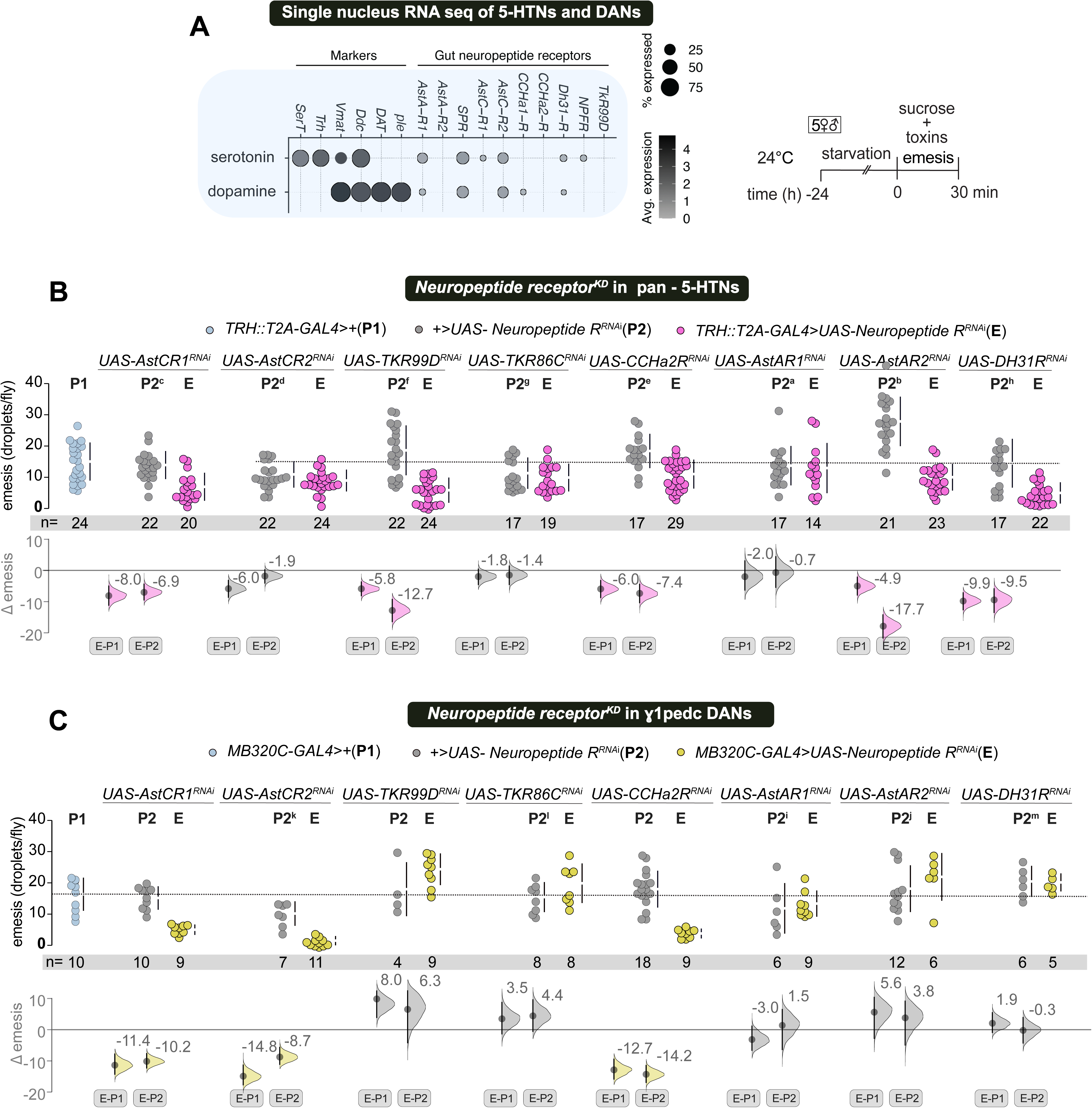
Neuropeptides released from the gut modulate 5-HTNs and DANs involved in emesis. **(A)** Single-cell transcriptomics data from (H. Li et al., 2022) show the expression of neuropeptide receptors in serotonergic neurons (5-HTNs) and dopaminergic neurons (DANs) involved in emesis. **(B)** Knockdown of *AstAR2*, *AstCR1*, *CCHa2R*, *TK99D*, and *DH31R* in *TRH::T2A-GAL4* reduces emesis at moderate to high levels compared to controls. **(C)** Knockdown of neuropeptide receptors in the mushroom body heel innervating, DANs labeled by *MB320C-GAL4* (PPL1-γ1pedc). *AstCR1*, *AstCR2*, and *CCHa2R* knockdown resulted in a very high reduction in emesis.

To test if the neuropeptides that are secreted from the gut signal to the brain 5-HTNs and DANs, we performed RNAi-mediated knockdown of their cognate receptors in the *TRH::T2A-GAL4* labeled 5-HTNs, and *MB320C-GAL4* labeled DANs. As a result, emesis was affected to varying degrees in the experimental groups (E), compared to the parental groups (P1: common GAL4/+ control and P2: the corresponding RNAi/+ controls) (Fig. 5B, C). For practical interpretation, we classified changes in emesis into three categories: none to low (mean effect sizes between 0 and 4), moderate (mean effect sizes between 4 and 8), and large (mean effect sizes over 8).

Knocking down *AstAR2*, *AstCR1*, *CCHa2R*, *TKR99D*, and *DH31R* in *TRH::T2A-GAL4* caused a moderate to large decrease in emesis (Fig. 5B; effect sizes highlighted with color). Therefore, their neuropeptide ligands (AstA, AstC, CCHa2, TK and DH31) contribute to emesis via 5-HTNs. Knockdown of *AstCR1*, *AstCR2*, and *CCHa2R* in *MB320C-GAL4* led to significant reduction in emesis (Fig. 5C). Using a broader *TH-GAL4* to target neuropeptide receptors confirmed our findings, with an additional strong effect observed with *AstAR1* (Fig. S5-1A). A 3-minute feeding test on 24-hour-starved flies showed no significant decrease in feeding across any of the lines tested (Fig. S5-1B, C). Our results suggest that toxin ingestion triggers the release of multiple neuropeptides from the gut, which then broadly signal to both 5-HTNs and DANs to promote emesis. Therefore, toxin-induced emesis is likely mediated by a complex interaction between neuropeptidergic and monoaminergic signaling pathways.

### Anticipatory emesis shares the 5-HT, DANS, and MB mediated pathway with naive emesis

The role of MB innervating PPL1 DANs in emesis (Fig. 4) and aversive learning in general (Aso et al., 2012; Das et al., 2014) suggests that flies might also learn from the aversive experience of ingesting toxins. Therefore, we designed an olfactory learning assay to explore whether flies can exhibit anticipatory emesis in expectation of toxin ingestion. Briefly, (Fig. 6A; see Materials and Methods for details), flies were grouped into small mixed-sex cohorts and exposed to a volatile odor for 30 minutes while ingesting a sugar-toxin mixture and exhibiting emesis. Subsequently, we tested 24-hour anticipatory emesis by exposing trained flies to the previously paired odor while allowing them to feed on blue-dyed sugar, and we counted any resulting emetic dots (experimental group). Similarly trained groups that were exposed to a novel odor while feeding on blue-dyed sugar served as controls. We calculated an anticipatory emesis index (AEI) by normalizing emesis droplets per fly in the experimental group to the average emetic droplets per fly in the control group (Fig. 6A: schema). We hypothesized that learning from the aversive toxin ingestion experience would cause the experimental groups to show higher anticipatory emesis than the controls (AEI > 1) (see Fig. S6-1A). Indeed, we observed anticipatory emesis when flies were trained with various emetic compounds, including copper sulfate, nicotine salts, caffeine, and lithium chloride. In contrast, no learning occurred with only 500 mM sucrose (Fig. 6B, See Supplementary video 3). We performed anticipatory emesis assays with multiple odor pairs to rule out the possibility that the phenotype was odor-specific. Based on our results, we typically used octan-3-ol (OCT) and methylcyclohexanol (MCH) odor pairs for all learning experiments (Fig. S6-1B). We also found that when trained with copper sulfate or a nicotine salt, anticipatory emesis decreased between 2 and 4 days after training (Fig. S6-1C).

**Figure 6.**
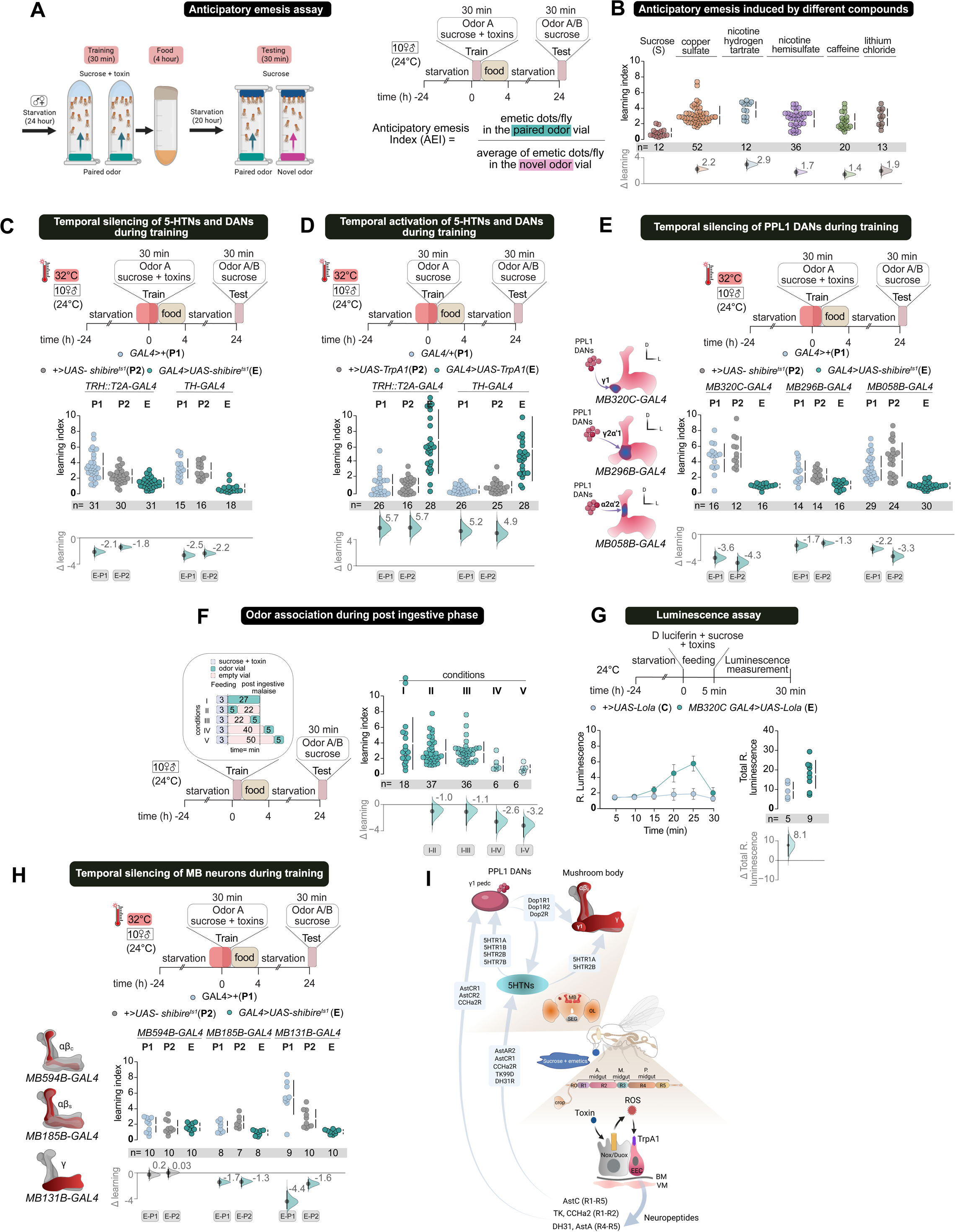
Anticipatory emesis recruits the same neural pathways as naive emesis. **(A)** Schematic for anticipatory emesis, Mixed sex flies are trained in the presence of a mixture of sugar and emetics in the presence of an odor for 30 min (left) and transferred to normal food for 4-5 hours. They are starved and tested after 24 hours of training (right). During testing one set of trained flies are allowed to feed on a mixture of sugar and blue dye, in the presence of the previously trained odor. Another set of trained flies are also fed on a mixture of sugar and blue dye, but in the presence of a novel odor for 30 mins. Anticipatory emesis index (AEI) is calculated by dividing the average of emetic dots/fly in the novel odor vial from emetic dots/fly in the paired odor vial. **(B)** Mixed sex flies exhibit anticipatory emesis in response to toxins and bitter-tasting compounds. **(C)** Transient silencing of 5-HTNs labeled by *TRH::T2A GAL4* and DANs labeled by *TH-GAL4* in flies using *UAS-Shibire^ts1^* during training impaired anticipatory emesis after 24 hours. **(D)** Transient artificial activation of 5HTNs labeled by *TRH::T2A GAL4* and DANs labeled by *TH-GAL4* using the *UAS-TrpA1* transgene induced anticipatory emesis in sugar fed flies. **(E)** Transient silencing PPL1-γ1pedc neurons (*MB320C-GAL4*), γ2ɑ’1 DANs (*MB296B-GAL4*) and ɑ2ɑ’2 (*MB058B-GAL4*) using *UAS-Shibire^ts1^* during training drastically reduced anticipatory emesis after 24 hours**. (F)** Odor association during the post ingestive phase induces anticipatory emesis. Flies were fed a sucrose–toxin mixture for 3 minutes and subsequently exposed to odor for different durations and intervals (left schema). Immediate post-feeding odor exposure, either for 27 minutes or 5 minutes (conditions I and II), induced robust anticipatory emesis 24 hours later. A 5-minute odor exposure given 22 minutes after feeding also elicited a comparable response (condition III). In contrast, little or no anticipatory emesis was observed when odor presentation was delayed by 40 minutes or more (conditions IV and V)**. (G)** γ1pedc PPL1 DANs exhibited persistent activity during post ingestive malaise phase. The luminescence activity of γ1pedc PPL1 DANs expressing *Lola-LUC* reporter (*MB320C GAL4>UAS-FLP;Lola>stop>LUC*) was recorded at six time points. These values were normalized by the mean luminescence of CS flies fed with nicotine (or sucrose) at each time point. The generated relative luminescence values for nicotine-fed and sucrose-fed groups was plotted. To calculate total relative luminescence, the average activity of individual flies across all six time points was determined for both *MB320C GAL4>UAS-FLP;Lola>stop>LUC* nicotine-fed and sucrose-fed groups. **(H)** Transient synaptic blockade of MB αβ-surface labeled by *MB185B-GAL4*, and MB γ-main neurons labeled by *MB131B-GAL4*, using *UAS-shibire^ts1^* during training impaired anticipatory emesis **(I)** Summary figure showing the aversive gut-brain axis underlying naive and anticipatory emesis uncovered by this work.

Like naive emesis, this anticipatory emesis also depends on neurotransmission from both *TRH::T2A-GAL4* labeled 5-HTNs and *TH-GAL4* labeled DANs (Fig. 6C). Artificial thermogenetic activation of these groups during training with only a sugar carrier caused increased anticipatory emesis, confirming that 5-HTNs and DANs serve as afferent, aversive teaching signals (Fig. 6D). Learning also relied on specific subsets of PPL1 DANs; reversible inhibition of γ1pedc (*MB320C-GAL4*), γ2α’1 (*MB296B-GAL4*), ɑ2ɑ’2 (*MB058B-GAL4*) groups reduced anticipatory emesis, suggesting their involvement in the learning process (Fig. 6E). Interestingly, blocking the latter two lines, which label γ2α’1 and α2α’2 DANs, had low to no effect on naive emesis. Similarly, silencing *TH-D4*, which includes PPL1, PPM2, and PPM3 neurons, reduces anticipatory emesis. We found no evidence for a role of PAM neurons labeled by *R58E02-GAL4* and *TH-C’-GAL4*, which includes the PPL2, PPM2, and PAL clusters in anticipatory emesis (Fig. S6-2A).Overall, we concluded that neural connections underlying naive and learned anticipatory emesis were shared.

### A post-ingestive malaise signal reinforces memory for anticipatory emesis

To explore the post-ingestive time window during which toxin sensation in the gut provides a teaching signal for learning, we performed an anticipatory emesis assay using *dTrpA1^1^* and flies with *dTrpA1* knocked down in EECs labeled by *R33A12-GAL4*. In both groups, anticipatory emesis was significantly reduced, indicating that the gut was the source of the teaching signal (Fig. S6-2B, C). We have demonstrated that ingesting toxins and detecting them in the midgut trigger emesis (Figs. 1, 2).

Consequently, we aimed to determine whether the same post-feeding process provides a teaching signal for anticipatory emesis. To investigate this, we allowed flies to feed on a sucrose and toxin mixture for 3 minutes in tubes. Afterwards, we transferred the flies to new tubes to expose them to OCT/MCH for varying durations.

In some cases, flies were not immediately exposed to an odor after feeding; instead, they were placed in odorless transition tubes between the feeding and odor exposure (Fig. 6F; schema). We observed that immediate odor exposure after feeding for both 27 minutes and 5 minutes induced anticipatory emesis 24 hours later (Fig. 6F; conditions I and II). Remarkably, when flies were exposed to an odor for 5 minutes, 22 minutes after feeding, they still showed an equally strong anticipatory emesis (Fig. 6F; condition III). However, little to no anticipatory emesis was observed when feeding and odor exposure were separated by 40 minutes or more (Fig. 6F; conditions IV and V). Our findings indicate that the teaching signal for anticipatory emesis occurs immediately after feeding and likely persists for approximately 20 to 40 minutes.

Since we identified a critical role for γ1pedc PPL1 DANs in both naive emesis and anticipatory emesis, and these neurons are known to reinforce the aversive valence of bitter food (Das et al., 2014), we then monitored their activity during the post-ingestive phase. To do this, we expressed a luminescence-based transcriptional reporter of neuronal activity, the immediate-early gene Lola promoter-driven luciferase transgene specifically in *MB320C-GAL4* (see Methods). Flies were fed sucrose and toxin mixture and then transferred to a luminometer to measure the post-feeding activity of the γ1pedc PPL1 DANs. Flies fed with toxin and sucrose showed a gradual increase in *MB320C-GAL4* activity, measured as relative luminescence, during the post-ingestive phase compared to flies fed sucrose alone. This signal peaked 25 minutes after feeding stopped and then declined, indicating a delayed and sustained activity of these neurons after feeding (Fig. 6G). The timing of γ1pedc PPL1 DAN activity overlaps with the post-ingestive time window in which malaise reinforced olfactory learning occurs (Fig. 6F), further confirming their role in emesis.

The MBs are the associative learning centers downstream to valence-specific reinforcement from the DANs, and therefore we tested their role in anticipatory emesis (Modi et al., 2020). We used MB lobe-specific split GAL4 lines to acutely inhibit synaptic transmission during training with *UAS-shibire^ts1^*. Our results indicate that neurotransmission from the MB αβ-surface (*MB185B-GAL4*) and the γ-main (*MB131B-GAL4*) neurons was essential for anticipatory emesis (Fig. 6H). These manipulations also prevented naive emesis (Fig.S6-2D). Our result suggests that MB neurons are integral to both naive emesis and anticipatory emesis and acts downstream of inputs from 5-HTNs and DANs.

Indeed, analysis of whole-brain single-cell transcriptomic data revealed that 5-HTRs and DARs are expressed in specific neuropils of the MBs (Croset et al., 2018) (Fig. S6-2E).

We knocked down the expression of 5-HTRs and DARs in the αβ-surface and γ-main neurons of the MB to assess their role in naive emesis (Fig. S6-2F-I). Knockdown of *Dop1R1* and *Dop1R2* in both MB lobes significantly reduced emesis (Fig. S6-2F, G).

The role of *Dop2R* in emesis was more prominent in the γ-main neurons than the αβ-surface neurons (Fig. S6-2G). Among the 5-HTRs, knocking down *5-HTR1A* in both lobes and *5-HTR2B* only in the γ-main compartment caused a moderate to considerable decrease in emesis (Fig. S6-2H, I). Therefore concluded that 5-HTRs and DARs are required in specific MB lobes to facilitate emesis.

## Discussion

This study identifies key endocrine and neural components of a gut-to-brain aversive interoceptive axis. Some of these components have clear mammalian counterparts. This axis conveys internal signals of sickness or malaise, especially after toxin ingestion, and initiates both naive emesis and notably the formation of an aversive memory for anticipatory emesis. We demonstrate that toxin exposure causes the release of a cocktail of neuropeptides from the midgut EECs, which act on central 5-HT and DA neurons located upstream of the MB neuropils. We also decipher a network composed of 5-HTNs, DANs, and MB neurons which coordinate both naive and anticipatory emesis.

### Emesis is triggered via the midgut

We used a high concentration of sucrose carrier to mask the toxins, to recreate situations where toxins are inadvertently ingested with food. Consequently, the toxin reached the gut undetected at the periphery, triggering emesis. This allowed us to study the underlying mechanisms and decipher the signaling network coordinating this response (Fig. 1, Fig. 6I). Our data suggest that a small amount of ingested toxins must initially reach the EECs in the midgut before emesis is triggered via robust crop contraction and likely involves the reverse action of at least the pump P1 in the proboscis (Supplementary video 2).

### Neuropeptide hormones represent the aversive afferent signal from the gut that drives emesis

We demonstrate that cationic dTrpA1 channels in the midgut EECs detect ROS produced in neighboring ECs due to toxin ingestion and then promote neuropeptide release (Fig. 2, Fig. 3). In mammals, irritants are similarly sensed by TrpA1 channels on secretory enterochromaffin cells in the small intestine, driving 5-HT release via a calcium-dependent mechanism (Bellono et al., 2017). Released 5-HT acts on 5-HT3 receptor-expressing vagal afferent nerve fibres that form synapse-like contacts with enterochromaffin cells in the intestinal villi (Bellono et al., 2017; Nozawa et al., 2009).

However, in this study, we found no evidence of 5-HT release from the fly gut, and the transcriptomics data of the midgut support this observation (data not shown). This is reminiscent of other endocrine pathways in flies and mammals, where different messengers have evolved to regulate similar behaviors and physiological processes. For instance, mammalian glucagon and insect adipokinetic hormone both control glucose and lipid homeostasis despite being evolutionary divergent (Y. Li et al., 2023).

Neuropeptide hormones have diverse functions depending on the specific state and neural circuit involved. Additionally, multiple neuropeptide hormones are known to be induced by the same stimuli. They may act on the same or different target organs or neural circuits to regulate similar or different downstream functions (Nässel, 2025). For example, the ingestion of amino acids triggers the release of multiple neuropeptides, including CCHa1, Tk, Dh31, and NPF from the gut. These neuropeptides mediate distinct processes after protein ingestion, such as sleep or arousal, suppression of protein appetite, promotion of sugar intake, and courtship by acting on neuronal targets both in the brain and in the gut (Ahrentløv et al., 2025; Malita et al., 2022; Titos et al., 2023; Yoshinari et al., 2024). Our systematic characterization of the pathways regulating emesis also identified a similar signaling motif where multiple gut neuropeptides target 5HTNs and DANs. Given the physiological risks of toxin ingestion, it is not surprising that multiple neuropeptides contribute to coordinating this protective response.

We report the release of five neuropeptides, AstA, AstC, CCHa2, Dh31, and Tk upon toxin ingestion. These neuropeptides probably coordinate different aspects of emesis by acting through the nervous system. In this study, we provide evidence for a gut-to-brain afferent axis, where gut neuropeptides transmit aversive sensory signals from ingested toxins to both 5-HTNs and the DANs that innervate the MB heels, primarily in combination. Additionally, gut-derived neuropeptides may locally influence the gut, associated neurons, and muscles to directly cause contractions of the crop and visceral muscles, regulate the opening or closing of sphincters, thus aiding emesis. Except for Tk, none of these peptides or their mammalian homologs has yet been identified as involved in emesis. Apart from 5-HT, mammalian ECs also express substance P (tachykinin/Tk) (Nässel & Zandawala, 2019; Zhong et al., 2021). Evidence suggests that gut-expressed Tk may influence emesis in mammals by affecting gut motility and the opening of the lower esophageal sphincter, which separates the stomach from the esophagus (Huber et al., 1993).

In a recent mouse study, staphylococcal toxin (SEA) induced 5-HT release from ECs, which signaled to Tac1+ (encoding substance P) excitatory neurons in the NTS of the DVC via afferent vagal neurons expressing the 5HT3 receptor. The DVC is considered the brain’s main gateway for interoceptive information from organs, including the gut. These Tac1+ DVC neurons ultimately mediated both retching-like mouth gaping and conditioned taste aversion learning (Xie et al., 2022). Similar functions for Tk are also observed in least shrews (Darmani et al., 2008). Here, we see Tk affects emesis via the TkR99D receptor in the 5-HTNs (Fig. 5). Hence, Tk peptides appear to play a conserved role in signaling aversion from the gut and inducing emesis in both flies and mammals.

### Serotonergic neurons convey an afferent gut-brain signal of toxin-induced distress from the gut

5-HT is a key neuromodulator involved in mammalian emesis, and many current antiemetic drugs target serotonergic signalling (Smith et al., 2012). Here, we have identified a broad serotonergic neuronal group, labelled by the *TRH::T2A-GAL4* (Fig. S4-1B), that plays a crucial role in vomiting. Multiple lines of evidence support a gut-to-MB afferent role for 5-HTNs in this cohort. A) All neuronal cell bodies in *TRH::T2A-GAL4* are located in the central brain (Fig. S4-1B), receiving emesis-relevant input from all the gut neuropeptides released upon toxin ingestion. B) All 5-HTRs are necessary in the MB-innervating DANs and the MBs themselves for emesis. C) Artificial stimulation of *TRH::T2A-GAL4* neurons reinforced an anticipatory emesis memory.

### Serotonergic neurons can influence crop contraction

Existing literature also indicates a role for 5-HTNs in brain-to-gut signaling that influences crop contraction. Although 5-HT is not produced by fly gut EECs, serotonergic innervation of the foregut occurs in many insect species, including flies (Budnik et al., 1989; Falibene et al., 2012; Moffett & Moffett, 2005; Molaei & Lange, 2003). In honeybees, 5-HT-immunoreactive neural processes innervate the esophagus, crop, proventriculus, and midgut. Further, 5-HTR transcripts are expressed in the crop and midgut. Inhibitors of 5-HTR inhibits contractions of the crop and proventriculus, indicating that excitatory 5-HT signaling regulates gut movement (French et al., 2014). Similar effects are observed in blowflies and fruit flies (Liscia et al., 2012; Solari et al., 2017). Consistent with this, a recent study shows that bitter taste activates serotonergic bitter-SELs in the fly brain, which project to the foregut-midgut junction and induce crop contraction via 5-HTR7 expressing enteric neurons. However, bitter-SELs driven crop contractions were reported not to cause regurgitation (Yao & Scott, 2022) Indeed, we find that inhibition of bitter-SELs activity does not impair emesis (Fig.4A, Fig. S4-1C). *TRH::T2A-GAL4* includes not only bitter-SELs but also additional neurons with axonal projections innervating the crop (Fig. S4-1B).

### Sustained, post-ingestive reinforcement of a delayed teaching signal from toxin ingestion

Like it does for other peripheral sensory aversive inputs, such as bitter taste and heat (Aso et al., 2012; Das et al., 2014; Galili et al., 2014), PPL1 DANs, especially the MB heel-innervating γ1pedc DANs, reinforce the sustained aversive signal from the gut caused by toxin ingestion. This is reflected by the persistent and heightened activity of γ1pedc DANs during this period following toxin ingestion, and explains how delayed associative learning can occur after toxin ingestion (Fig. 6F, G). Such a persistent signal could be due to prolonged release of neuropeptides from EECS, triggered by continuing sensing of elevated ROS levels in the gut (Fig. 2).

### Limitations of this study

In this study we were unable to isolate the subset of *TRH::T2A-GAL4* neurons that receive afferent signals from the gut and communicate with the MB-innervating DANs in inducing naive and anticipatory emesis. Consequently, we were unable to provide evidence for a direct neural connection between the *TRH::T2A-GAL4* labeled 5-HTNs and the MB-innervating DANs. Additionally, we did not determine the role of crop-innervating and anterior midgut innervating neurons labeled by *TRH::T2A-GAL4* in emesis. These experiments must await generation of reagents to target specific subsets of 5-HTNs, as is available for DAN populations. Crop contraction *per se* does not ensure emesis. Strong contractions of the crop has to be coordinated with the closing of the proventricular sphincter, the opening of the foregut sphincters, and the reverse activity of the cibarial and esophageal pumps for effective emesis (Stoffolano & Haselton, 2013). This complete emetic sequence in the gut is probably synchronized by 5-HT and other yet to be identified pathways.

### Future perspectives

*Drosophila* emerges from our work as the only genetically tractable model organism capable of recapitulating the molecular and physiological pathways of naive emesis. Importantly our model of learned anticipatory emesis allows us to model chemotherapy induced anticipatory vomiting in flies, and potentially offering an opportunity to screen specific drugs against this process. The components of the aversive gut-brain axis that we have uncovered may also be involved in transmitting gut-derived interoceptive cues during acute or chronic infections, injuries, or drug-induced damages. Emerging research in both flies and mammals suggests that some neurodegenerative diseases may originate in the gut and then spread to the brain (Loh et al., 2024). The aversive gut signaling identified here may also play a crucial role in this process.

## Materials and Methods

### **A.** Fly rearing

Flies were reared on standard laboratory food at 23- 25 ℃ and 60-70% humidity, with a 12 :12 h light::dark cycle inside incubators. All the experiments were performed in (Wild type Canton-S; CS) flies. For a list of transgenic fly stock genotypes and sources, see supplementary table 1.

#### Chemicals and other Reagents

See Supplementary table 2.

### **B.** Behavior assays

#### i) Emesis assay

The behavioral setup includes three custom-made, transparent, circular acrylic sheets. The central sheet is designed with 12 circular arenas, each measuring 35 mm in diameter and 1.7 mm in height. Uniform illumination is provided by an LED light source placed beneath the setup to ensure consistent backlighting across all arenas.

Behavioral activity is recorded, when needed, from above using a Logitech HD webcam (1080p resolution, 30 frames per second), mounted at a fixed height to capture all arenas simultaneously with minimal distortion.

The stock solution of all the emetic compounds mentioned was freshly prepared in RO water. The desired concentrations were achieved by diluting stock solutions and mixing them with 750 mM sucrose, 0.8% Brilliant blue dye, and 0.75% agar. The final volume of each mixture was adjusted using RO water.

Adult *Drosophila melanogaster* (5–6 days old) were food-deprived on moist filter paper for approximately 23–24 hours. After deprivation, a mixed group of five flies was placed into each arena. Flies were given 30–40 µL of a test mixture containing sucrose and the emetic compound inside the arena, and allowed to feed *freely* for 30 minutes. At the end of this period, the assay was stopped, and arenas were immediately transferred to a – 80 °C freezer to euthanize the flies. Emetic droplets deposited within each arena were then counted and recorded manually.

#### **ii)** Anticipatory emesis assay

The stock solution for all the emetic compounds was freshly prepared in RO water. The desired concentrations were achieved by diluting stock solutions and mixing them with 500 mM sucrose, 0.8% Brilliant blue dye, and 0.75% agar. The final volume of each preparation was adjusted with RO water.

0.5–0.8 µL of octan-3-ol (OCT) or 0.5–1 µL of 4-methylcyclohexanol (MCH) was diluted in 1.25 mL of white oil. This was considered approximately a 1000-fold dilution. For odor presentation, 20 µL of the diluted odorant was applied onto a piece of Whatman filter paper, which was then placed on the inside of a 5 mL tube lid. The tube was sealed with a piece of parafilm punctured with small holes to allow for odor diffusion.

A mixed population of 5 to 6-day-old flies was starved on 0.75% agar for 24 hours before training. During training, flies were transferred to pre-odorized 5 mL plastic tubes containing a food mixture of 500 mM sucrose and emetics in 0.75% agar. They were allowed to feed in this environment for 30 minutes. Afterward, flies were moved to fresh 5 mL tubes with non-toxic food for 4-5 hours, then starved again for up to 24 hours. For testing, trained flies were divided into two groups: one was transferred to 5 mL tubes pre-odorized with the trained odor, and the other group to tubes pre-odorized with a novel odor. In both cases, flies were allowed to feed on a mixture of 500 mM sucrose and 0.8% blue dye in 0.75% agar for 30 minutes. After testing, flies were sacrificed, and the blue dots were counted manually.

#### **iii)** PER assay

24-hour food-deprived wild-type flies (5-6 days old) were anesthetized for 1 minute by placing them in a cold test tube immersed in a 4°C ice bath. The flies were then immobilized on their backs on a glass slide using a non-toxic adhesive and left to recover for 1 hour at 25°C with 65% relative humidity. These flies were initially offered water on their legs to identify responders, which were then used for subsequent experiments. The responders were satiated with water and tested with sample solutions applied to their forelegs. Compounds were added to 750 mM sucrose, and the effect on proboscis extension was examined. Each fly received the same solution three times with water applications in between, and proboscis extensions were recorded. Different flies were used to test different substances, while the same flies were used to test various concentrations of the same solution, starting with the lowest concentration.

#### **iv)** Latency assay

Mixed-sex groups of flies, with about five flies per group, were starved for 24 hours, transferred to arenas, and allowed to feed on emetic food (750 mM sucrose, toxin, and blue dye in 0.75% agar). Their behavior was videotaped for 40 minutes and analyzed. The time to the first emetic episode (first droplet deposition) from the time the flies were introduced into the arena was manually annotated from the recordings for each fly.

#### **v)** Tracking Defecation

Flies were treated the same as those for the latency assay described above, but their behavior was video-recorded for 120 minutes. The timing of individual defecation events was manually annotated from the recorded videos.

#### **vi)** Pharmacological screen

To evaluate the effects of serotonin and dopamine receptor antagonists, as well as antioxidant treatment, various pharmacological agents were incorporated into 0.75% agar. Different concentrations of S1A WAY (a 5-HT1A receptor antagonist), ketanserin (a 5-HT2 receptor antagonist), SB258719 (a 5-HT7 receptor antagonist), flupentixol (a D1 receptor antagonist), and dithiothreitol (DTT, an antioxidant) were used. Flies were starved for 24 hours on this agar. After starvation, the flies underwent an emesis assay as described above.

#### **vii)** Thermogenetic experiments

*UAS- shibire^ts1^*and *UAS-dTrpA1* flies were raised at 20°C- 22 °C and transferred to 32°C 30 minutes before the experiments to induce transgene expression.

#### **viii)** Belly-mount preparation for quantifying crop contraction imaging

The belly-mount preparation was adapted from (Koyama et al., 2020) with modifications. Wild-type flies were starved for 24 hours before the experiment. Flies were anesthetized with a brief 20-second burst of CO₂, and their dorsal side, mainly the wings, was affixed to an imaging glass slide using clear nail polish. To achieve optimal positioning and reduce interference during crop visualization, the hind legs were removed. A small rectangle of clear Elmer’s Liquid School Glue (Amazon, B06WVDBR62), about the size of the fly’s abdomen, was applied to the center of a 40-mm coverslip and carefully placed on the ventral side of the fly’s abdomen (Fig. 1E). Fine adjustments were made to ensure clear visualization of the internal organs. Flies were allowed to habituate in this state for 2–3 minutes before proceeding. They were then fed a mixture of sucrose and toxin or sucrose alone. Crop contractions were manually counted through a stereomicroscope, and videos were recorded simultaneously for later analysis.

### **C.** Feeding quantification

Mixed-sex groups of flies, each consisting of approximately five flies, were starved for 24 hours, transferred to arenas, and allowed to feed on emetic food for 5 minutes. The feeding duration was selected because all flies in the arenas feed sufficiently within that time, which also corresponds with the typical latency before they begin to vomit.

Flies were then transferred to 1.5 ml tubes and homogenized in PBS. The homogenate was centrifuged, and the supernatant was collected. The absorbance of the supernatant was measured at 630 nm (for Brilliant Blue FCF dye) using a spectrophotometer to quantify the amount of ingested dyed food.

### **D.** Immunohistochemistry

The adult brains and guts were gently dissected in phosphate-buffered saline (PBS) using forceps and then fixed in 4% formaldehyde in PBS for 1 hour at room temperature. After three washes in PBST (PBS with 0.3% Triton X-100), tissues were blocked in 5% normal goat serum in PBST for 1 hr at room temperature. The tissues were then incubated with primary antibodies in a blocking solution at 4 °C for 2 days. After three washes in PBST, the tissues were incubated overnight at 4 °C in a blocking solution with secondary antibodies. The tissues were then washed in PBST, followed by one to two exchanges of PBS, and mounted on poly-L-lysine-coated slides. Images were captured using an inverted Zeiss LSM 880 confocal microscope at 10x and 20x magnification.

### **E.** ROS staining

ROS levels in live tissue were measured as previously described (Owusu-Ansah et al., 2008). Flies starved for 20–24 hours were fed a mixture of sucrose, toxin, and agar for 10 minutes in a vial. Guts were then dissected and incubated with 30 mM DHE for 7 minutes at 25°C in the dark. After four washes with PBS, the guts were mounted and immediately imaged for red fluorescence signals using an inverted Zeiss LSM 880 confocal microscope at 40x and 60x magnification with fixed gain and offset settings for both experimental and control sugar-fed samples.

### **F.** Luminescence assay

The luminescence assay for detecting neuronal activity in freely behaving adult flies was adapted from (Guo et al., 2017). Briefly, flies were starved for 24 hours before the assay and then transferred into 5 mL Eppendorf tubes containing a) 5% sucrose, emetic, and 20 mM D-luciferin in 2% low-melting agarose. Control flies were placed in tubes with the same mixture, but importantly, without the emetic. Flies were allowed to feed for 5 minutes and subsequently transferred to a white, flat-bottom 96-well plate, with one fly per well. The flies were loaded through cross-shaped cuts over individual wells in the film covering the plate. The flaps prevented flies from escaping. The plate was loaded into a Promega Luminometer, and the luminescence signal was recorded every 5 min over a 35 min period to coincide with the post-ingestive phase of toxin ingestion.

Luminescence values from the experimental group (wild-type × Lola-luciferase) under both feeding conditions (toxin + sucrose or sucrose alone) were averaged across biological replicates at each time point. These values were used to normalize the luminescence signals from the GAL4 × Lola-luciferase flies under the corresponding feeding conditions (N. luminescence). The cumulative sum of N. luminescence values across time points was plotted as total relative luminescence.

### **G.** Single cell/nucleus-transcriptome analyses

Single-nucleus transcriptomes for the gut, 5HTNs and DANs were generated previously (Li et al., 2022). Single-cell RNA transcriptomes for KCs were generated previously (Croset et al., 2018). All analyses were conducted with Seurat (v 4.4.0) for R Studio (2024.04.2+764).

### **H.** Connectome analyses

Neurons known to express 5-HT were catalogued previously (Eckstein et al., 2024). See the database here: https://github.com/funkelab/drosophila_neurotransmitters for further details. We used the most-stringent confidence interval of 5 (i.e.evidence for protein expression in the given celltype) to subsets of 5-HTNs for analysis. The morphology of these neurons matched the neurons covered by *TRH::T2A-GAL4*. 5-HTN and PPL1 neuron reconstructions were downloaded using FAFBseg (v 0.15.3) and cloud-volume library (v 12.4.1, https://github.com/seung-lab/cloud-volume) through reticulate (v 1.43.0) and python (v 3.10.16) in R Studio (2025.05.2 Build 522), and visualized using RGL (v 1.3.24). We used FlyWire v783 for all analyses. Synapses (Princeton table; Date: 24 Jul 2025) and cell annotations (Date: 14 May 2025) were retrieved from Codex (Yu et al., 2025). Influence scores were calculated using influencer v 0.1.0 (https://github.com/natverse/influencer) based on the connectome influence calculator (Ajabi et al., 2025). We used a lenient const c of 24 for adjusted influence that corresponds to the minimum accepted influence of 3.78e-11 (Bates et al., 2025). Root ids of neurons used for the analyses are provided- Supplementary table 3.

### **I.** Data plotting and statistics

Statistical analysis was performed as previously described (Thakare et al., 2024). Briefly, GraphPad Prism version 8.3.1 or the website: www.estimationstats.com (Ho et al., 2019) was used to plot the data and generate all the estimation/delta plots. All data are represented as scatter plots, with error bars indicating the mean and a 95% confidence interval (CI). The effect size is calculated as the mean difference between the two groups. This represents the magnitude of the difference between the two groups. The actual effect size is shown as the dot in the spread. The two black lines that extend from the effect size dot denote the 95% CI and represent the precision of the effect size. The CI of the effect size was generated by 5000 bias-corrected and accelerated (BCa) bootstrap resampling.

## Acknowledgments

We thank the Bloomington Drosophila Stock Center and the Vienna Drosophila Resource Center for fly stocks. We thank Dr. Sheeba Vasu, JNCASR, and Dr. Girish R., IISER Pune, for generously sharing reagents and fly lines. We thank Dr. Dick Nässel for providing feedback on the manuscript and Dr. Alexander Bates for sharing the influencer package prior to its public release. We also extend our gratitude to the members of the Brain and Feeding Behavior Lab for their constructive feedback and constant support. Also, a special thank you to all the trainees who had worked on this project, especially Pavithra US, Aditi Manjare, Vaikheri, Nikita. RK received a doctoral fellowship from the University Grants Commission (UGC). Both RK and MS are currently supported by a Pratiksha Trust Extra-Mural Support for Transformational Aging Brain Research grant (EMSTAR/2023/SL03) awarded to GD. PC received a doctoral fellowship from the University Grants Commission (UGC) and was briefly supported by a Ramanujan Fellowship award to GD (SB/S2/RJN-048/2017). NB is funded by the University Grants Commission (UGC). This project was initially funded by extramural funding from an erstwhile SERB (now ANRF) grant (CRG/2019/005587). The GD lab is also funded by extramural funds from the Pratiksha Trust Extra-Mural Support for Transformational Aging Brain Research grant (EMSTAR/2023/SL03) and the Department of Biotechnology, Government of India grant (BT/PR51490/MED/ 22/361/2024). The lab also receives generous intramural funding support from BRIC-NCCS, Pune. Schematics were created with the help of <u>Biorender.com</u>.

## Author contributions

G.D. designed the emesis assay and its refinements with input from R.K. and P.C. The anticipatory emesis assay was developed by G.D. and R.K., with R.K. further developing it. R.K. performed all anticipatory emesis assays. The luciferase assay and the Bellymount assay were adapted and modified by R.K. Dissections, immunostaining, and confocal imaging were mainly carried out by R.K. and M.S. Naive emesis assays were conducted by R.K., P.C., N.B., and M.S. Z.C. and M.Z. curated and plotted all the single-cell transcriptomics data and all fly brain connectomics data. R.K., P.C., N.B., M.S., and Z.C. performed all experiments, data analysis, and plotting for the manuscript. M.Z. and G.D. supervised the project. G.D. secured funding and directed the research. R.K. and P.C. wrote the initial drafts of the manuscript. G.D. primarily wrote the final manuscript with input from R.K., M.Z., and N.B.

## Abbreviations

DVC: Dorsal vagal complex
NTS: Nucleus tractus solitarius
AP: Area postrema
5-HT: Serotonin
DA: Dopamine
SP: Substance P
DMV: Dorsal motor nucleus of the vagus
ROS: Reactive oxygen species
EECs: Enteroendocrine cells
ECs: enterocytes
5-HTNs: Serotonergic neurons
DANs: Dopaminergic neurons
DARs: Dopamine receptors
5-HTRs: Serotonin receptors
MBs: Mushroom bodies
PER: Proboscis extension response
Gr: gustatory receptor
GrNs: gustatory receptor neurons
DPM: Dorsal paired medial neurons
PAM: Protocerebral anterior medial neurons
AEI: Anticipatory emesis index
OCT: Octan-3-ol
MCH: Methylcyclohexanol
MBONs: Mushroom body output neurons
NHT: nicotine hydrogen tartrate
NHS: nicotine hydrogen sulfate
DTT: dithiothreitol
ANV: Anticipatory nausea and vomiting
*Gene*/Protein: Gene names
*SerT*: Serotonin transporter
*TH*/TH: Pale/tyrosine hydroxylase
*Vmat*/Vmat: Vesicular monoamine transporter
*TrpA1*/TrpA1: Transient receptor potential ankyrin 1
*Nox*/Nox: NADPH oxidase
*Duox*/Duox: Dual oxidase
*Trh/* Trh: Tryptophan hydroxylase
*Trhn*/Trhn: Tryptophan hydroxylase neuronal
*AstA*/AstA: Allatostatin A
*AstC*/AstC: Allatostatin C
*Tk*/Tk: Tachykinin
*CCHa2*/CCHa2: CCHamide-2
*Dh31*/Dh31: Diuretic hormone 31
*amon*/Amon: amontillado.

