## Supplemental Data File 1 for "A gut-brain axis for aversive interoception drives innate and anticipatory emesis in *Drosophila*"

### Supplementary Materials

### Figure:S1-1

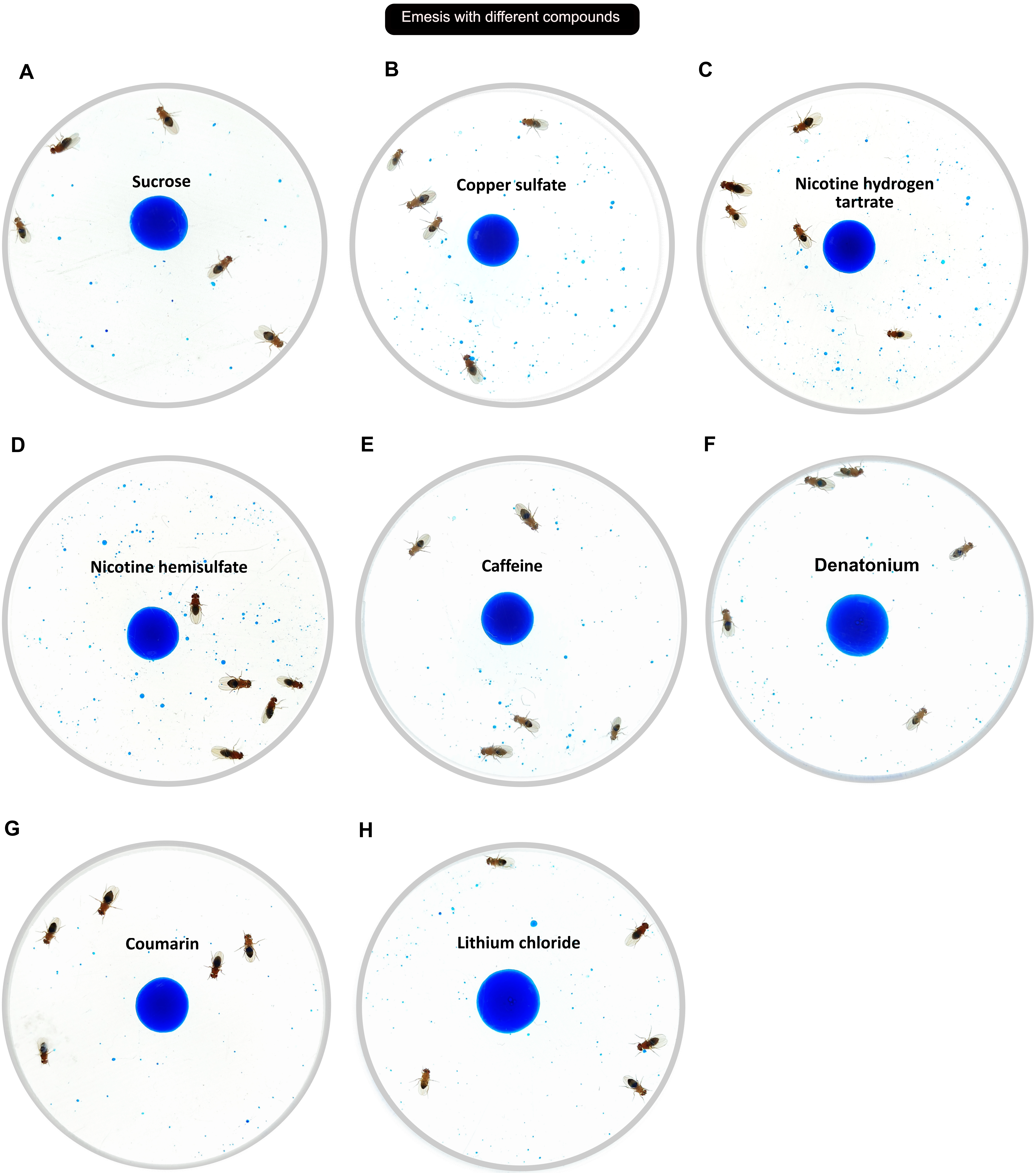

Emesis with different compounds
A
Sucrose
B
Copper sulfate
C
Nicotine hydrogen tartrate
D
Nicotine hemisulfate
E
Caffeine
F
Denatonium
G
Coumarin
H
Lithium chloride

**Figure S1-1. Emesis exhibited due to the ingestion of various emetic compounds.** Emesis observed after 30 min with 5 flies, after the ingestion of emetics (Fig. 1B). Representative arena images are shown. **(A)** 750 mM carrier sucrose without emetics. Emesis to 750 mM carrier sucrose mixed with: **(B)** copper sulfate, **(C)** nicotine hydrogen tartrate, **(D)** nicotine hemisulfate, **(E)** caffeine, **(F)** denatonium, **(G)**, coumarin, and **(H)** lithium chloride.

### Figure: S1-2

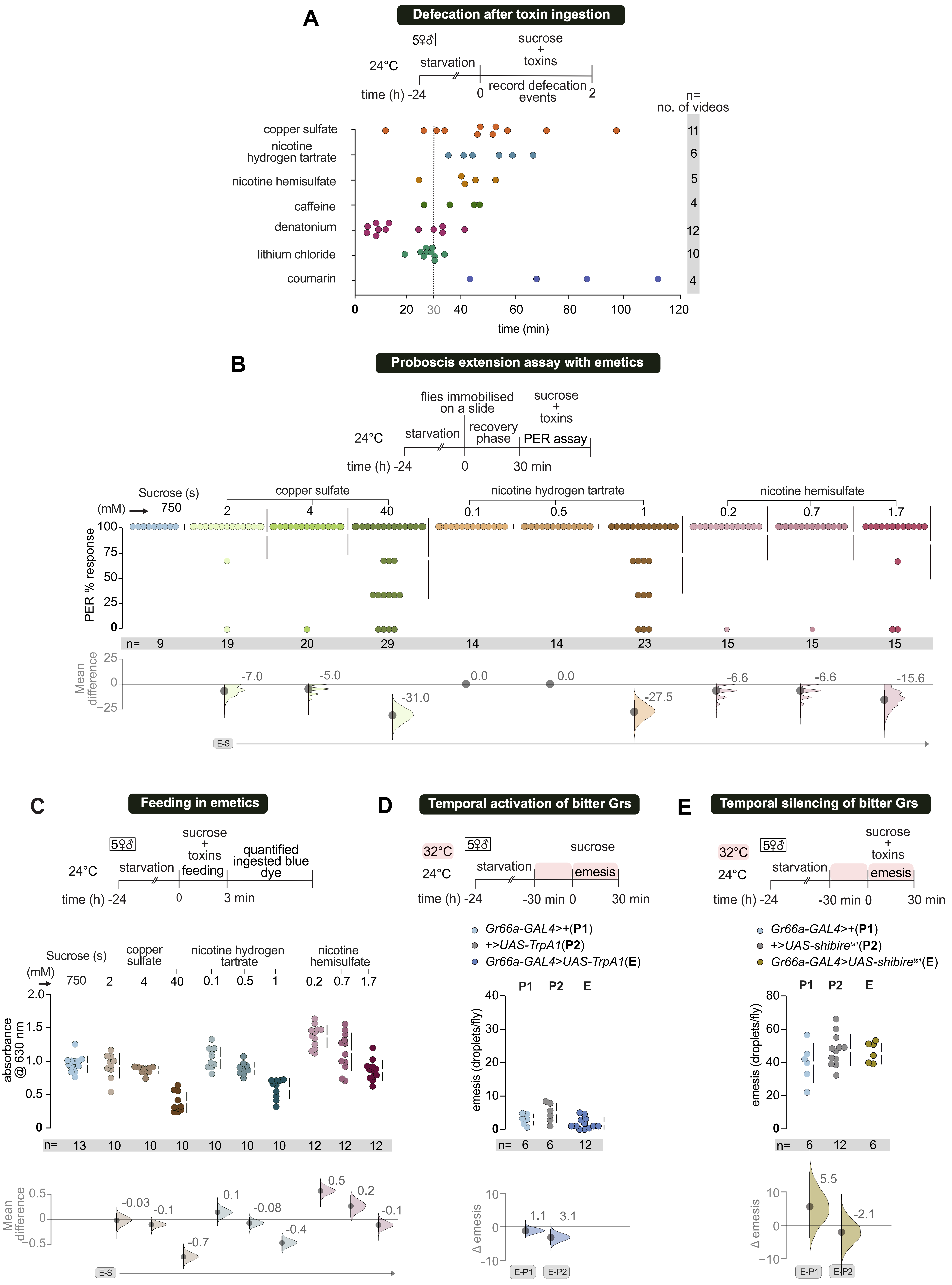

A
Defecation after toxin ingestion
5♀♂
24°C
starvation
sucrose + toxins
record defecation events
time (h) -24
0
2
n= no. of videos
copper sulfate
nicotine hydrogen tartrate
nicotine hemisulfate
caffeine
denatonium
lithium chloride
coumarin
11
6
5
4
12
10
4
time (min)
B
Proboscis extension assay with emetics
flies immobilised on a slide
sucrose + toxins
starvation
recovery phase
PER assay
30 min
Sucrose (s)
copper sulfate
nicotine hydrogen tartrate
nicotine hemisulfate
(mM)
PER % response
n=
Mean difference
E-S
C
Feeding in emetics
feeding
quantified ingested blue dye
3 min
absorbance @ 630 nm
D
Temporal activation of bitter Grs
32°C
sucrose
emesis
-30 min 0
Gr66a-GAL4>+(P1)
+>UAS-TrpA1(P2)
Gr66a-GAL4>UAS-TrpA1(E)
emesis (droplets/fly)
Δ emesis
E-P1
E-P2
E
Temporal silencing of bitter Grs
+>UAS-shibire ts1 (P2)
Gr66a-GAL4>UAS-shibire ts1 (E)

**Figure S1-2. Involvement of aversive taste, feeding, crop function and defecation in emesis.**

**(A)** Defecation is absent during the 30-minute emesis assay for most of the compounds tested, except denatonium. Flies in arenas with constant access to sugar-toxin mixture were video recorded for 2 hours. The recordings were monitored and analyzed for defecation events. **(B)** Suppression of Proboscis extension response (PER) by varying toxin concentrations. Hardly any PER suppression is seen at the $e^{max}$ concentration of the compounds used. Higher concentrations of toxins visibly suppress PER, indicating sensory detection of bitter compounds. **(C)** Feeding quantification by blue dye ingestion for sugar + toxins used in the previous panel. Consistent with PER suppression, feeding was suppressed at higher than $e^{max}$ concentration of toxins. **(D)** Acute thermogenetic activation of bitter gustatory receptor neurons (GrNs) does not induce emesis. Flies expressing *UAS-TrpA1* in *GR66a-GAL4* labeled bitter-sensing neurons were placed at 32 °C to and fed 750 mM sucrose alone, but did not exhibit increased emesis compared to parental controls. **(E)** Transient inhibition of bitter GrNs labeled by *GR66a-GAL4* neurons by the expression of *UAS-Shibire*$^{ts1}$ transgene and raising the flies to 32 °C did not inhibit emesis.

### Figure S2-1

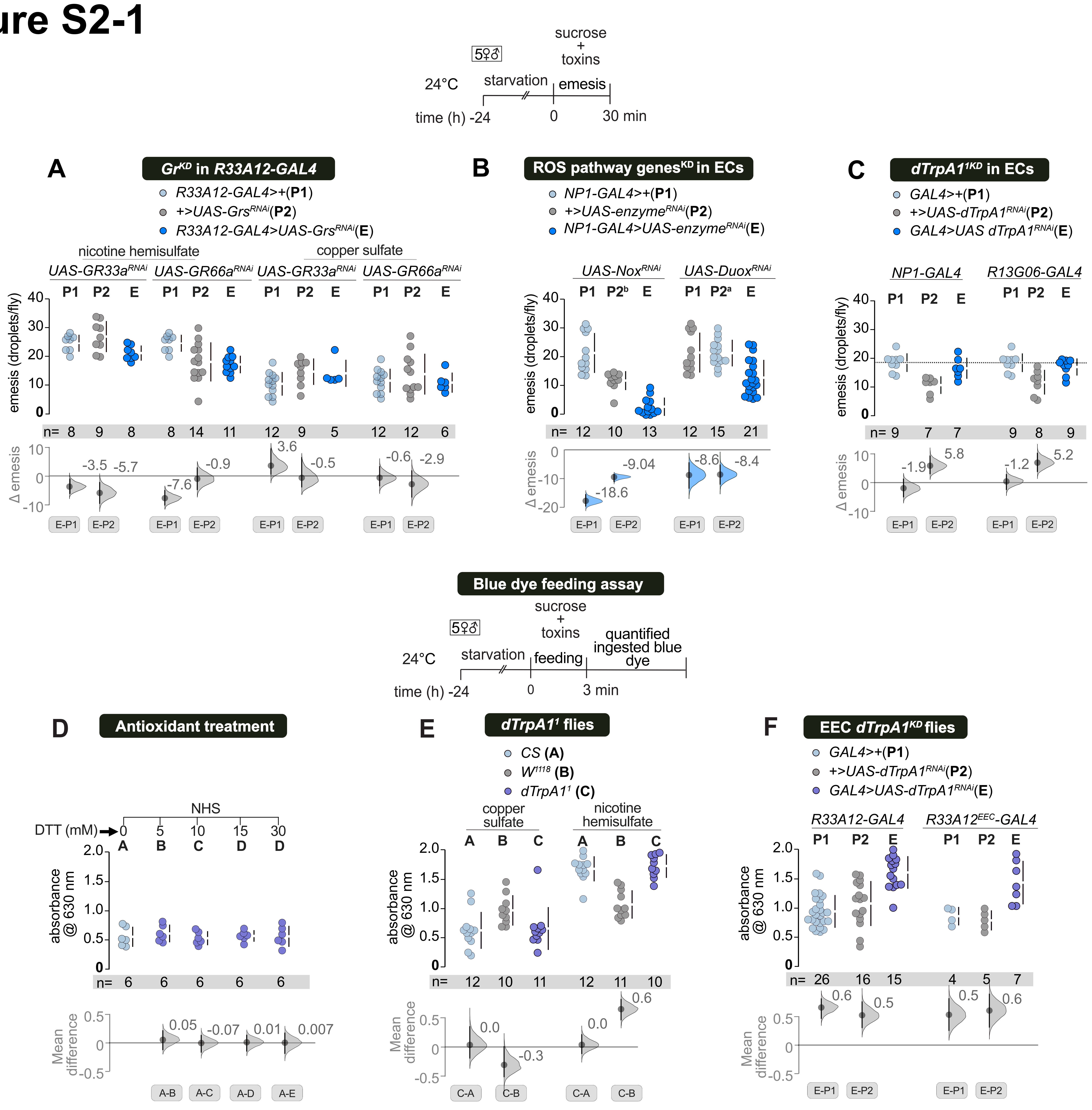

sucrose + toxins
24°C
starvation
emesis
time (h) -24
0
30 min
A
Gr^KD in R33A12-GAL4
R33A12-GAL4>+(P1)
+>UAS-Grs^RNAi(P2)
R33A12-GAL4>UAS-Grs^RNAi(E)
nicotine hemisulfate
copper sulfate
UAS-GR33a^RNAi
UAS-GR66a^RNAi
emesis (droplets/fly)
Δ emesis
B
ROS pathway genes^KD in ECs
NP1-GAL4>+(P1)
+>UAS-enzyme^RNAi(P2)
NP1-GAL4>UAS-enzyme^RNAi(E)
UAS-Nox^RNAi
UAS-Duox^RNAi
C
dTrpA1^1KD in ECs
GAL4>+(P1)
+>UAS-dTrpA1^RNAi(P2)
GAL4>UAS dTrpA1^RNAi(E)
NP1-GAL4
R13G06-GAL4
Blue dye feeding assay
feeding
quantified ingested blue dye
3 min
D
Antioxidant treatment
NHS
DTT (mM)
absorbance @ 630 nm
Mean difference
E
dTrpA1^1 flies
CS (A)
W^1118 (B)
dTrpA1^1 (C)
F
EEC dTrpA1^KD flies
GAL4>+(P1)
+>UAS-dTrpA1^RNAi(P2)
GAL4>UAS-dTrpA1^RNAi(E)
R33A12-GAL4
R33A12^EEC-GAL4

**Figure S2-1. ROS-producing genes in EECs, but not bitter GRNs, mediate emesis. (A)** Knockdown of bitter *GrNs* in EECs does not impair emesis. Knockdown of *GR33a* and *GR66a* in EECs labeled by *R33A12-GAL4* does not affect emesis. **(B)** Knockdown of ROS-inducing genes in EECs reduces emesis. UAS-RNAi knockdown of *NOX* and *DUOX* in ECs using *NP1-GAL4* inhibits emesis. **(C)** RNAi-mediated knockdown of *dTrpA1* in ECs labeled by *NP1-GAL4* and *R13G06-GAL4* does not impair emesis. **(D)** Short-term feeding assay with blue-dye ingestion in flies preloaded with different DTT concentrations shows no reduction compared to the control. **(E)** $dTrpA1^{1}$ flies display feeding behavior similar to control wild-type strains like *CS* and $w^{1118}$ when fed with carrier sugar, emetics, and blue dye. **(F)** Knockdown of *dTrpA1* in *R33A12-GAL4* and $R33A12\text{-}GAL4^{EEC}$ does not impair feeding in flies. On the contrary feeding is increased robustly in the experimental flies.

### Figure S2-2

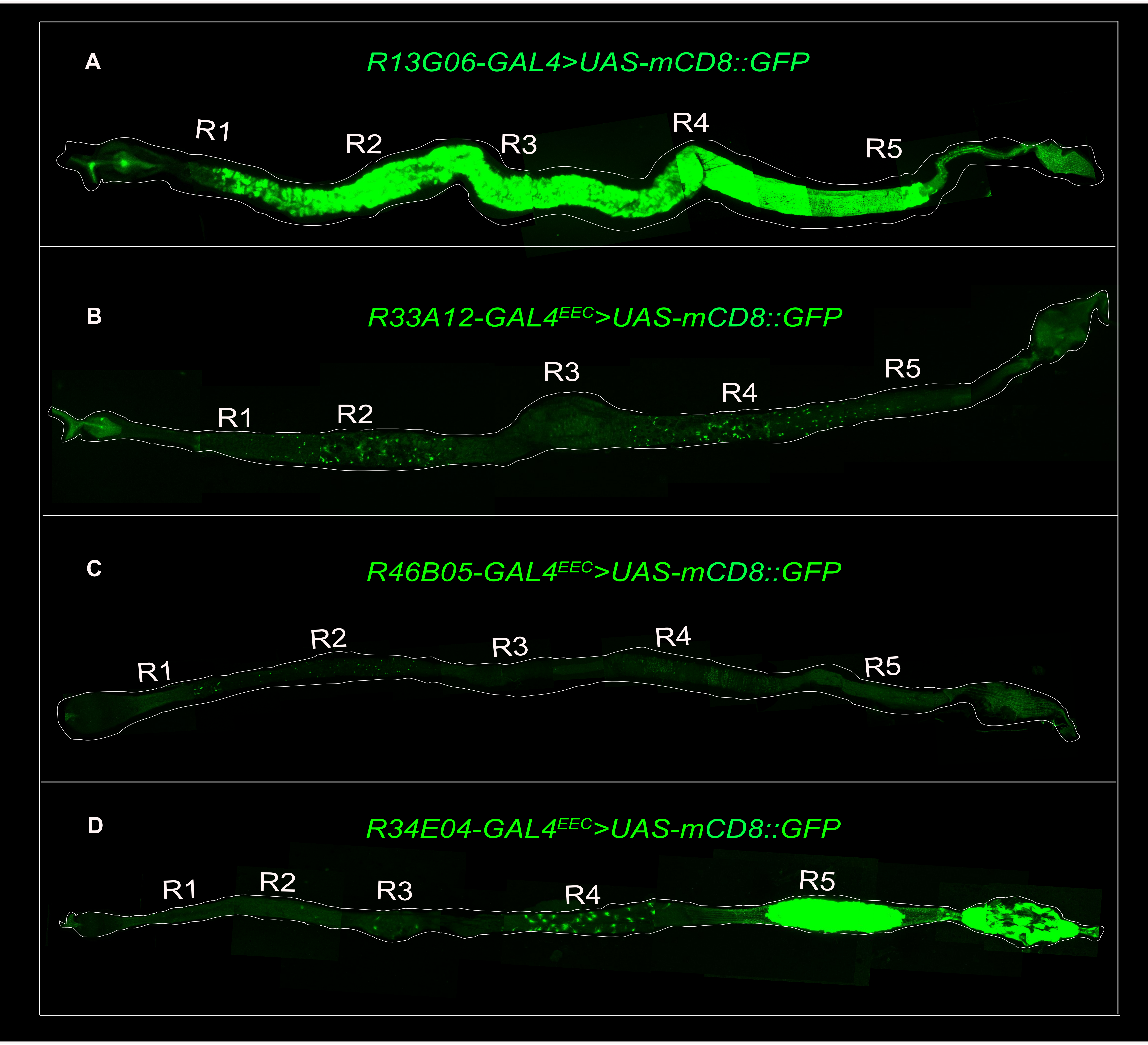

A
R13G06-GAL4>UAS-mCD8::GFP
R1
R2
R3
R4
R5
B
R33A12-GAL4EEC>UAS-mCD8::GFP
R1
R2
R3
R4
R5
C
R46B05-GAL4EEC>UAS-mCD8::GFP
R1
R2
R3
R4
R5
D
R34E04-GAL4EEC>UAS-mCD8::GFP
R1
R2
R3
R4
R5

**Figure S2-2. GAL4 driver lines with expression in ECs and EECs.**

GAL4 lines labeled using *UAS-mCD8::GFP* **(A)** *R13G06-GAL4* broadly label ECs in the fly gut **(B)** *R33A12 GAL4*$^{EEC}$ labels broad EEC in the fly gut (R1-R5). **(C)** *R46B05-GAL4*$^{EEC}$ labels EECs in the anterior midgut (R1-R3) **(D)** *R34E04-GAL4*$^{EEC}$ labels EECs in the posterior midgut (R4-R5).

### Figure:S3-1

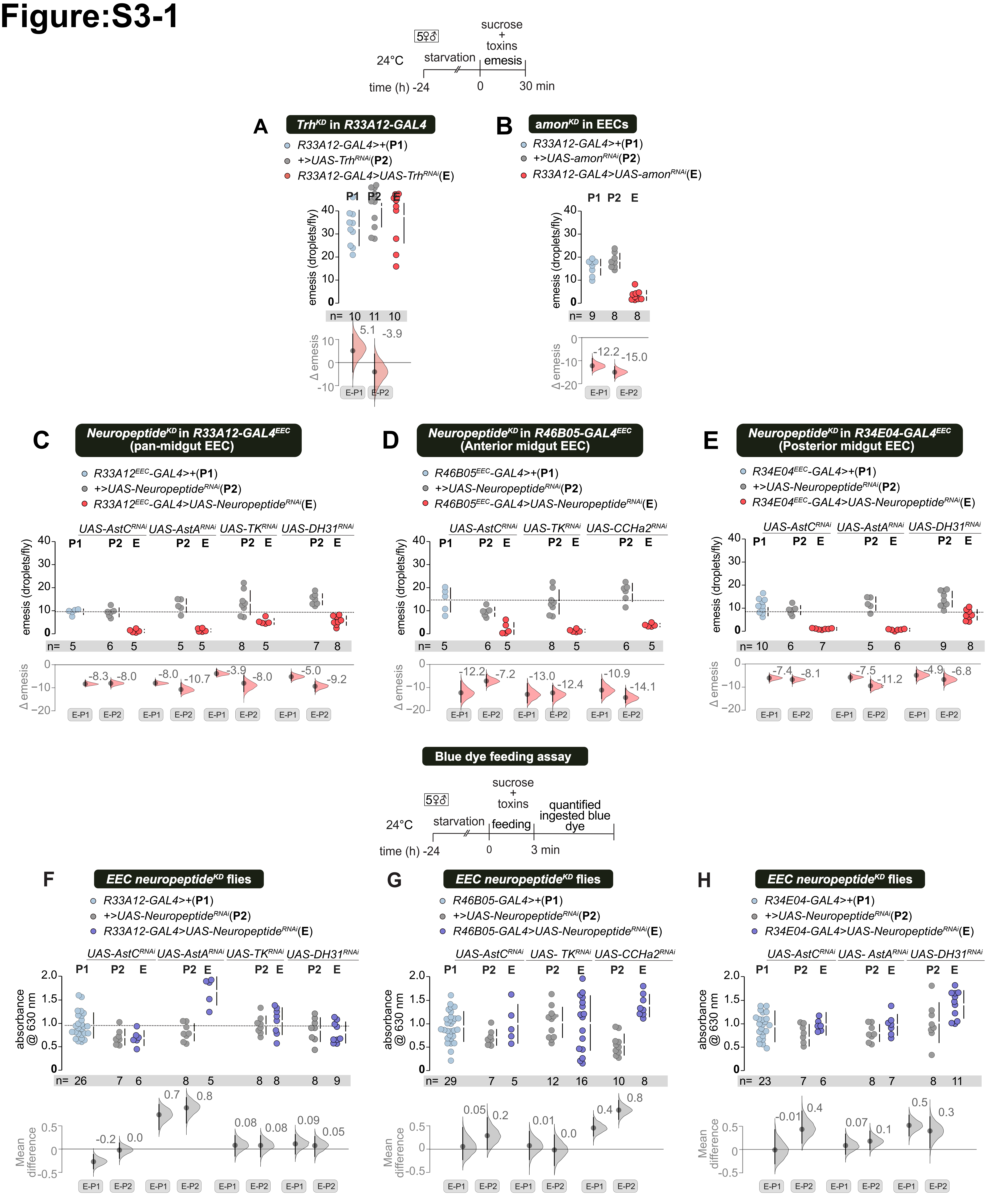

24°C
5♀♂
starvation
sucrose + toxins emesis
time (h) -24
0
30 min
A
Trh^KD in R33A12-GAL4
R33A12-GAL4>+(P1)
+>UAS-Trh^RNAi(P2)
R33A12-GAL4>UAS-Trh^RNAi(E)
B
amon^KD in EECs
R33A12-GAL4>+(P1)
+>UAS-amon^RNAi(P2)
R33A12-GAL4>UAS-amon^RNAi(E)
emesis (droplets/fly)
Δ emesis
C
Neuropeptide^KD in R33A12-GAL4^EEC (pan-midgut EEC)
R33A12^EEC-GAL4>+(P1)
+>UAS-Neuropeptide^RNAi(P2)
R33A12^EEC-GAL4>UAS-Neuropeptide^RNAi(E)
UAS-AstC^RNAi
UAS-AstA^RNAi
UAS-TK^RNAi
UAS-DH31^RNAi
D
Neuropeptide^KD in R46B05-GAL4^EEC (Anterior midgut EEC)
R46B05^EEC-GAL4>+(P1)
+>UAS-Neuropeptide^RNAi(P2)
R46B05^EEC-GAL4>UAS-Neuropeptide^RNAi(E)
UAS-AstC^RNAi
UAS-TK^RNAi
UAS-CCHa2^RNAi
E
Neuropeptide^KD in R34E04-GAL4^EEC (Posterior midgut EEC)
R34E04^EEC-GAL4>+(P1)
+>UAS-Neuropeptide^RNAi(P2)
R34E04^EEC-GAL4>UAS-Neuropeptide^RNAi(E)
UAS-AstC^RNAi
UAS-AstA^RNAi
UAS-DH31^RNAi
Blue dye feeding assay
24°C
5♀♂
starvation
sucrose + toxins feeding
quantified ingested blue dye
time (h) -24
0
3 min
F
EEC neuropeptide^KD flies
R33A12-GAL4>+(P1)
+>UAS-Neuropeptide^RNAi(P2)
R33A12-GAL4>UAS-Neuropeptide^RNAi(E)
G
EEC neuropeptide^KD flies
R46B05-GAL4>+(P1)
+>UAS-Neuropeptide^RNAi(P2)
R46B05-GAL4>UAS-Neuropeptide^RNAi(E)
H
EEC neuropeptide^KD flies
R34E04-GAL4>+(P1)
+>UAS-Neuropeptide^RNAi(P2)
R34E04-GAL4>UAS-Neuropeptide^RNAi(E)
absorbance @ 630 nm
Mean difference

**Figure S3-1. Neuropeptide expression, but not 5HT expression from the midgut EECs, is necessary for emesis.**

**(A)** Knockdown of tryptophan hydroxylase (*Trh*), in the midgut EEC labeled by *R33A12-GAL4*, does not impair emesis **(B)** Inhibition of *amon*, the *Drosophila* prohormone convertase 2, in the midgut EECs with *R33A12-GAL4* almost completely abolishes emesis. **(C, D, E)** Identified neuropeptides that facilitate emesis (identified in Fig. 3) were inhibited with the same *UAS-RNAi* lines in *R33A12-GAL4*$^{EEC}$, *R46B05-GAL4*$^{EEC}$, and *R34E04-GAL4*$^{EEC}$, which are EEC-specific versions of the corresponding GAL4 lines created by combining them with *nSyb-GAL80*. All showed reduced emesis, further confirming our results. **(F, G, H)** Short-term feeding assay in flies was fed sugar + emetic + blue dye, corresponding to positive hits in (Fig. 3B - D). No decrease in feeding was observed; in fact, increased feeding was seen in some cases compared to parental controls. The dashed line in the graphs indicates the mean of the common control group P1.

### Figure: S4-1

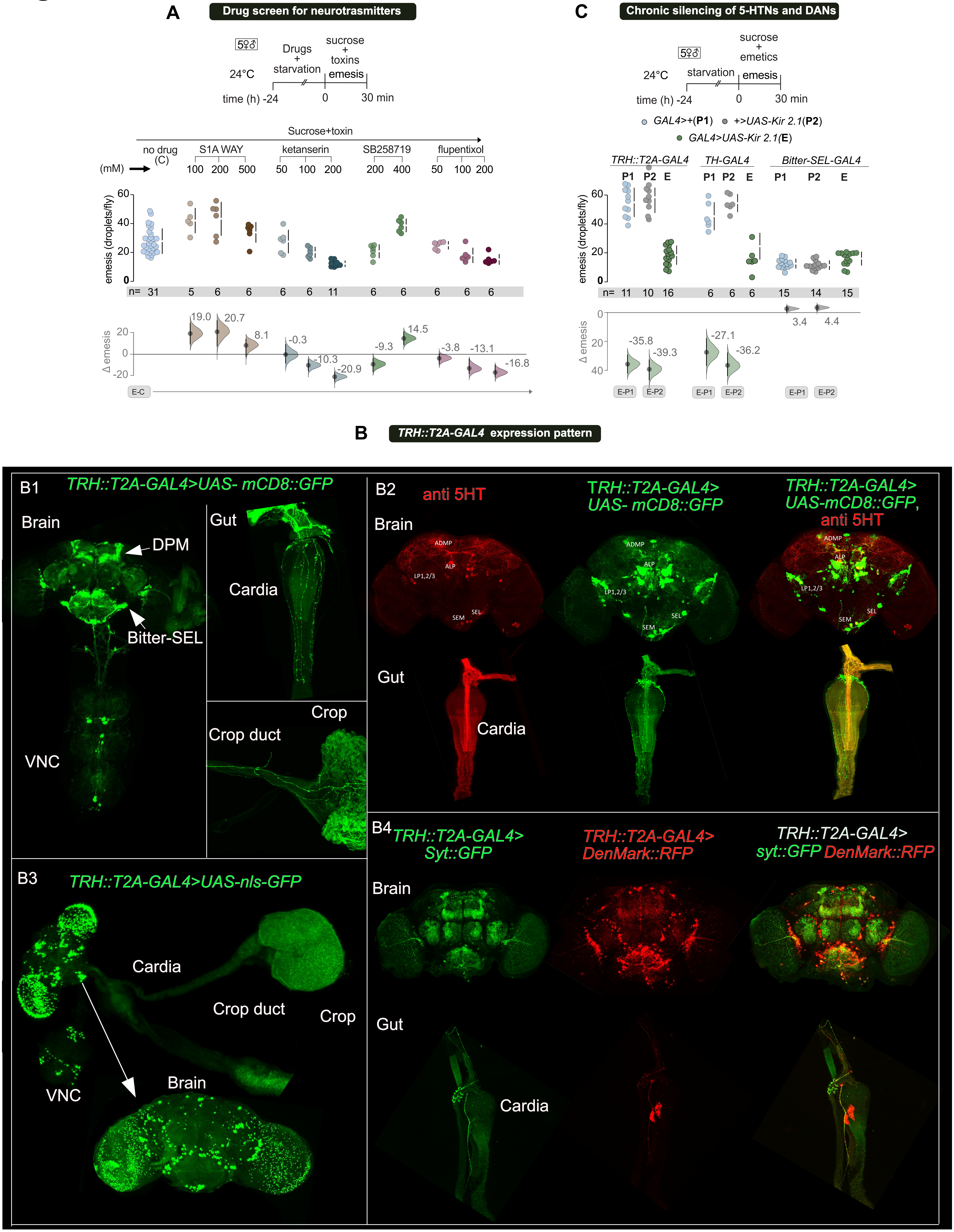

A
Drug screen for neurotrasmitters
5♀3♂
24°C
Drugs + starvation
sucrose + toxins
emesis
time (h) -24
0
30 min
Sucrose+toxin
no drug (C)
S1A WAY
ketanserin
SB258719
flupentixol
(mM)
100 200 500
50 100 200
200 400
50 100 200
emesis (droplets/fly)
n= 31 5 6 6 6 6 11 6 6 6 6 6
19.0
20.7
8.1
-0.3
-10.3
-20.9
-9.3
14.5
-3.8
-13.1
-16.8
Δ emesis
E-C
C
Chronic silencing of 5-HTNs and DANs
5♀3♂
24°C
starvation
sucrose + emetics
emesis
time (h) -24
0
30 min
GAL4>+(P1)
+>UAS-Kir 2.1(P2)
GAL4>UAS-Kir 2.1(E)
TRH::T2A-GAL4
TH-GAL4
Bitter-SEL-GAL4
P1 P2 E
n= 11 10 16 6 6 6 15 14 15
-35.8
-39.3
-27.1
-36.2
3.4
4.4
E-P1
E-P2
B
TRH::T2A-GAL4 expression pattern
B1
TRH::T2A-GAL4>UAS- mCD8::GFP
Brain
DPM
Bitter-SEL
VNC
Gut
Cardia
Crop
Crop duct
B2
anti 5HT
TRH::T2A-GAL4> UAS- mCD8::GFP
TRH::T2A-GAL4> UAS-mCD8::GFP, anti 5HT
Brain
ADMP
ALP
LP1,2/3
SEM
SEL
Gut
Cardia
B3
TRH::T2A-GAL4>UAS-nls-GFP
Cardia
Crop duct
Crop
VNC
Brain
B4
TRH::T2A-GAL4> Syt::GFP
TRH::T2A-GAL4> DenMark::RFP
TRH::T2A-GAL4> syt::GFP DenMark::RFP
Brain
Gut
Cardia

**Figure S4-1. 5-HTNs and DANs modulate emesis**

**(A)**. Known dopamine and serotonin receptors (DARs and 5-HTRs) inhibitors modulated emesis. **(B)** **B1.***TRH::T2A-GAL4* broadly labels serotonergic neurons in the brain, ventral nerve cord (VNC), and gut. **B2**.Co-labelling of *TRH::T2A-GAL4* expression with anti *5-HT* antibody **B3**. *UAS-nls-GFP* expression of *TRH::T2A-GAL4*. **B4.** Axonal and dendritic projections of *TRH::T2A-GAL4* expressing neurons. *Syt::GFP* (green) labels presynaptic terminals, and *DenMark::RFP* (red) marks dendritic compartments. **(C)** Chronic silencing of 5-HTNs and DANs reduces emesis. Disruption of 5-HTNs labeled by *TRH::T2A- GAL4* and DANs labeled by *TH-GAL4* using *UAS-Kir2.1* impaired emesis. However, *Bitter-SEL-GAL4* had no effect.

### Figure: S4-2

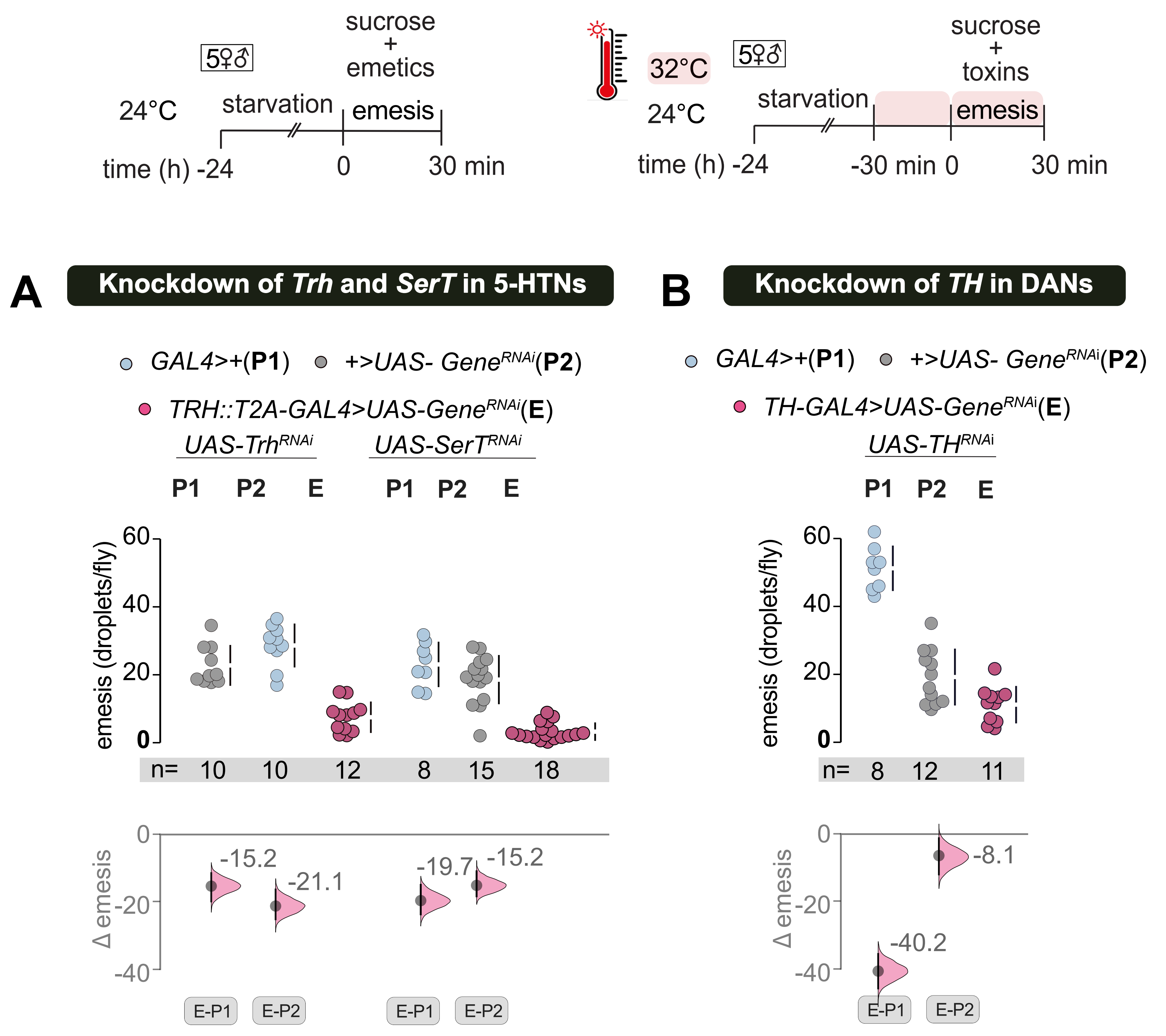


C

Temporal silencing of aversive DANs

|  | PAM | PAL | PPM1 | PPM2 | PPM3 | PPL1 | PPL2 |
|---|---|---|---|---|---|---|---|
| anti-TH | ~100 | 4.9±0.1 | 0.8±0.1 | 7.9±0.3 | 6.8±0.2 | 11.7±0.2 | 5.5±0.2 |
| TH | 8.5±1.0 | 5.0±0 | 4.0±0 | 8.3±0.25 | 7.0±1.0 | 11.0±0 | 6.0±0 |
| TH-C' | 0.3±0.2 | 3.0±0.1 | 0.5±0.1 | 5.4±0.2 | 0 | 0 | 5.9±0.1 |
| TH-D' | 0.1±0.1 | 1.3±0.1 | 0 | 2.7±0.3 | 5.8±0.3 | 9.8±0.1 | 0 |
| TH-D4 | 0 | 0 | 0 | 1.6±0.1 | 3.4±0.2 | 5.6±0.1 | 0 |

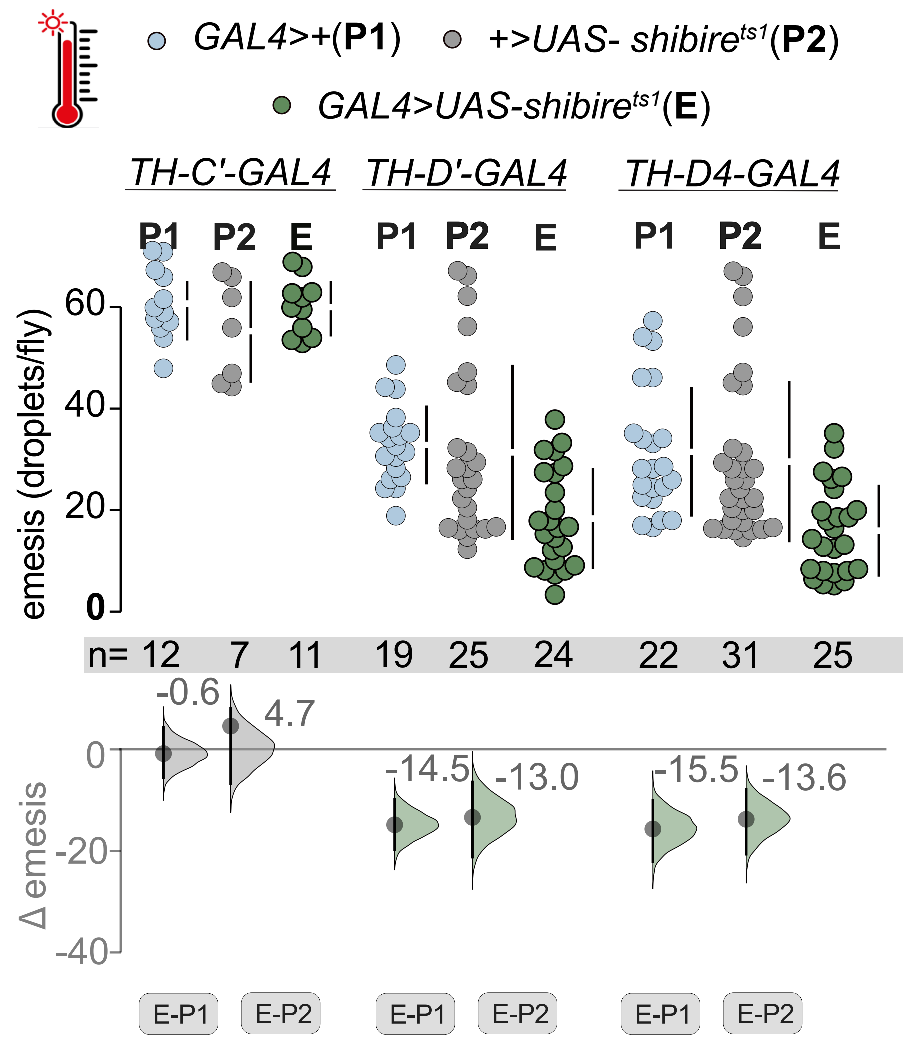


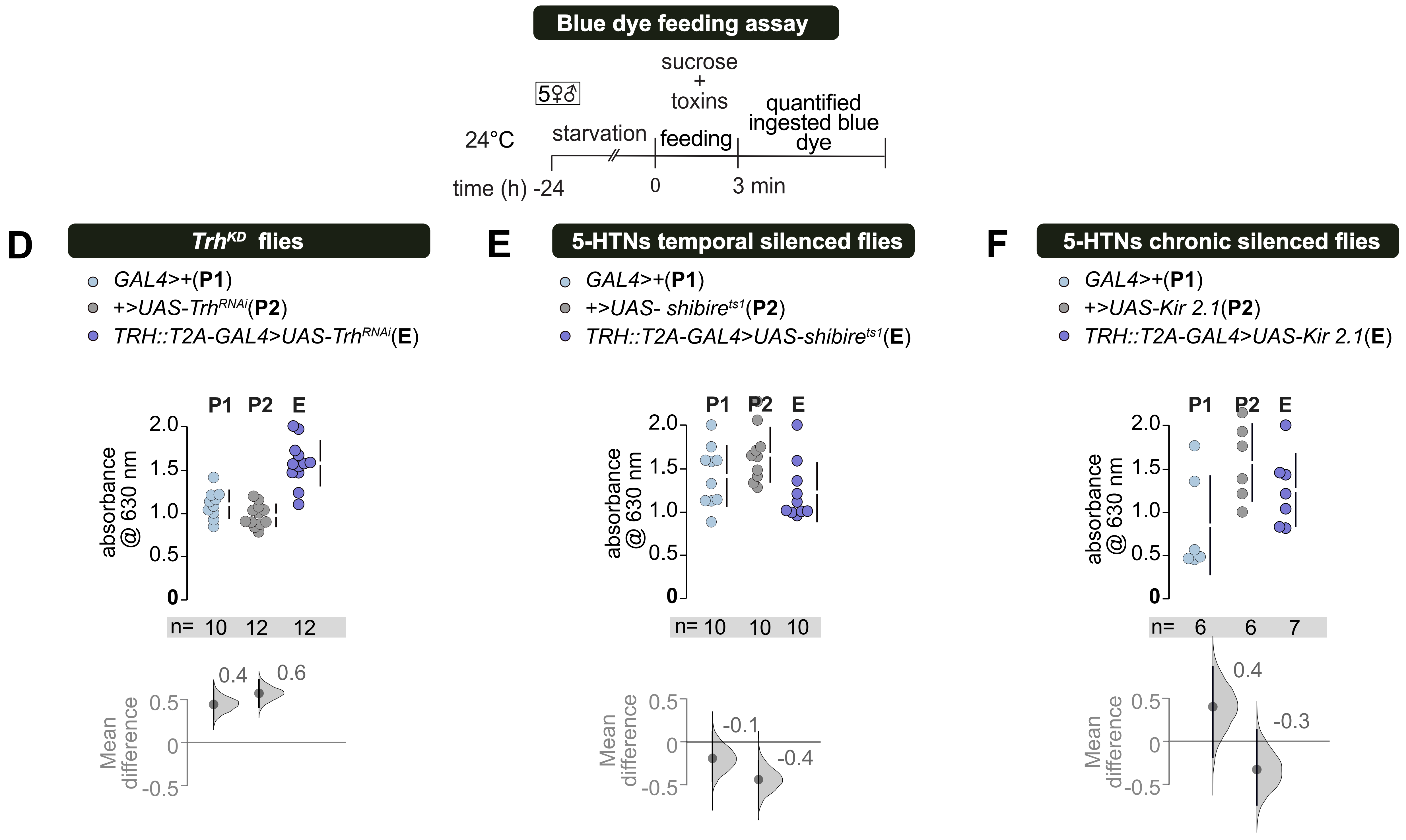

**Figure S4-2. Knockdown of 5-HTRs in TH-D4-GAL4 labeled dopaminergic neurons reduces emesis.**

**(A)** Knockdown of *Trh* and *SerT* in 5-HTNs labeled by TRH::T2A- GAL4 impaired emesis. **(B)** Knockdown of *tyrosine hydroxylase* (*TH*) in DANs labeled by *TH-GAL4* reduced emesis. **(C)** PPL1 DANs are likely needed for emesis. *TH*+ cells labeled by *nls-GFP* driven by each *Gal4* line, as determined by immunostaining experiments with anti-GFP and anti-*TH* antibodies modified from (Liu et al., 2012, S1-Table) (table) Genetic screen for identifying the emesis relevant sub-cluster of *TH-GAL4* labeled DANs reveals that transient, thermogenetic silencing of *TH-D'-GAL4* and *TH-D4-GAL4*, which label PPL1, PPM2, and PPM3 clusters, reduced emesis, while silencing of *TH-C'-GAL4*, which excludes PPL1 neurons had no effect. **(D)** Knockdown of *Trh* in 5-HTNs labeled by *TRH::T2A-GAL4* does not reduce short-term feeding. **(E)** Silencing 5-HTNs labeled by *TRH::T2A-GAL4* using *UAS-Shibire*$^{ts1}$ does not alter feeding**. (F)** Silencing 5-HTNs labeled by *TRH::T2A-GAL4* using *UAS-Kir2.1* does not impair feeding.

### Figure: S4-3

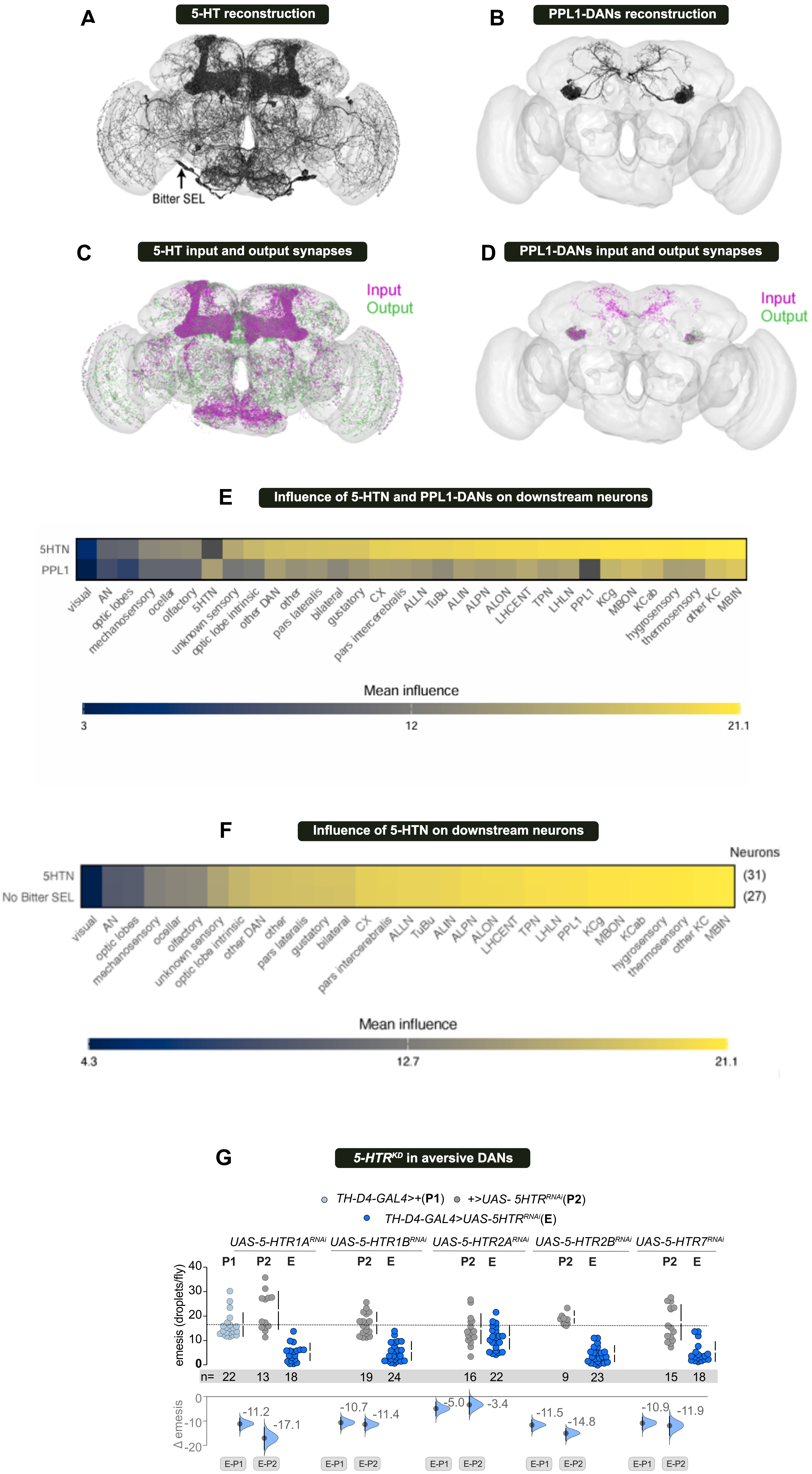
A
5-HT reconstruction
Bitter SEL
B
PPL1-DANs reconstruction
C
5-HT input and output synapses
Input
Output
D
PPL1-DANs input and output synapses
Input
Output
E
Influence of 5-HTN and PPL1-DANs on downstream neurons
5HTN
PPL1
visual
AN
optic lobes
mechanosensory
ocellar
olfactory
5HTN
unknown sensory
optic lobe intrinsic
other DAN
other
pars lateralis
bilateral
gustatory
CX
pars intercerebralis
ALLN
TuBu
ALIN
ALPN
ALON
LHCENT
TPN
LHLN
PPL1
KCg
MBON
KCab
hygrosensory
thermosensory
other KC
MBIN
Mean influence
3
12
21.1
F
Influence of 5-HTN on downstream neurons
Neurons
5HTN
No Bitter SEL
(31)
(27)
visual
AN
optic lobes
mechanosensory
ocellar
olfactory
unknown sensory
optic lobe intrinsic
other DAN
other
pars lateralis
gustatory
bilateral
CX
pars intercerebralis
ALLN
TuBu
ALIN
ALPN
ALON
LHCENT
TPN
LHLN
PPL1
KCg
MBON
KCab
hygrosensory
thermosensory
other KC
MBIN
Mean influence
4.3
12.7
21.1
G
5-HTR^KD in aversive DANs
TH-D4-GAL4>+(P1)
+>UAS- 5HTR^RNAi(P2)
TH-D4-GAL4>UAS-5HTR^RNAi(E)
UAS-5-HTR1A^RNAi
UAS-5-HTR1B^RNAi
UAS-5-HTR2A^RNAi
UAS-5-HTR2B^RNAi
UAS-5-HTR7^RNAi
P1 P2 E
P2 E
P2 E
P2 E
P2 E
emesis (droplets/fly)
40
30
20
10
0
n= 22 13 18 19 24 16 22 9 23 15 18
Δ emesis
0
-10
-20
-11.2
-17.1
-10.7
-11.4
-5.0
-3.4
-11.5
-14.8
-10.9
-11.9
E-P1
E-P2
E-P1
E-P2
E-P1
E-P2
E-P1
E-P2
E-P1
E-P2

**Figure S4-3. Interactions between 5-HTNs and PPL1 DANs**

**(A)** Reconstructions of neurons in the FlyWire connectome (Dorkenwald et al., 2024; Schlegel et al., 2024) that are known to express 5-HT based on experimental evidence (Eckstein et al. 2024, https://github.com/funkelab/drosophila_neurotransmitters). Bitter-SELs (Yao & Scott, 2022) have been indicated with an arrow. **(B)** Reconstructions of PPL1 DANs subtypes implicated in emesis. Input (magenta) and output (green) synapses of **(C)** 5-HTNs and **(D)** PPL1 neurons. **(E)** Influence of 5-HTN and PPL1 activation on different neuronal classes. The number of starter neurons activated are indicated on the right. Neuron classes implicated in emesis are shaded in orange. Influence of starter neurons on themselves is excluded (dark grey square). **(F)** Activation of 5-HTNs or 5-HTNs excluding bitter-SELs has similar influence on downstream neuronal classes. **(G)** RNAi-mediated knockdown of fly serotonin receptors (5-HTRs), *5-HTR1A*, *5-HTR1B*, *5-HTR2B*, and *5-HTR7*, in the broad dopaminergic line *TH-D4-GAL4* (includes PPL1, PPM2, PPM3), shows a considerable reduction in emesis. The effect is low to moderate when *5-HTR2A* is knocked down.

### Figure: S5-1

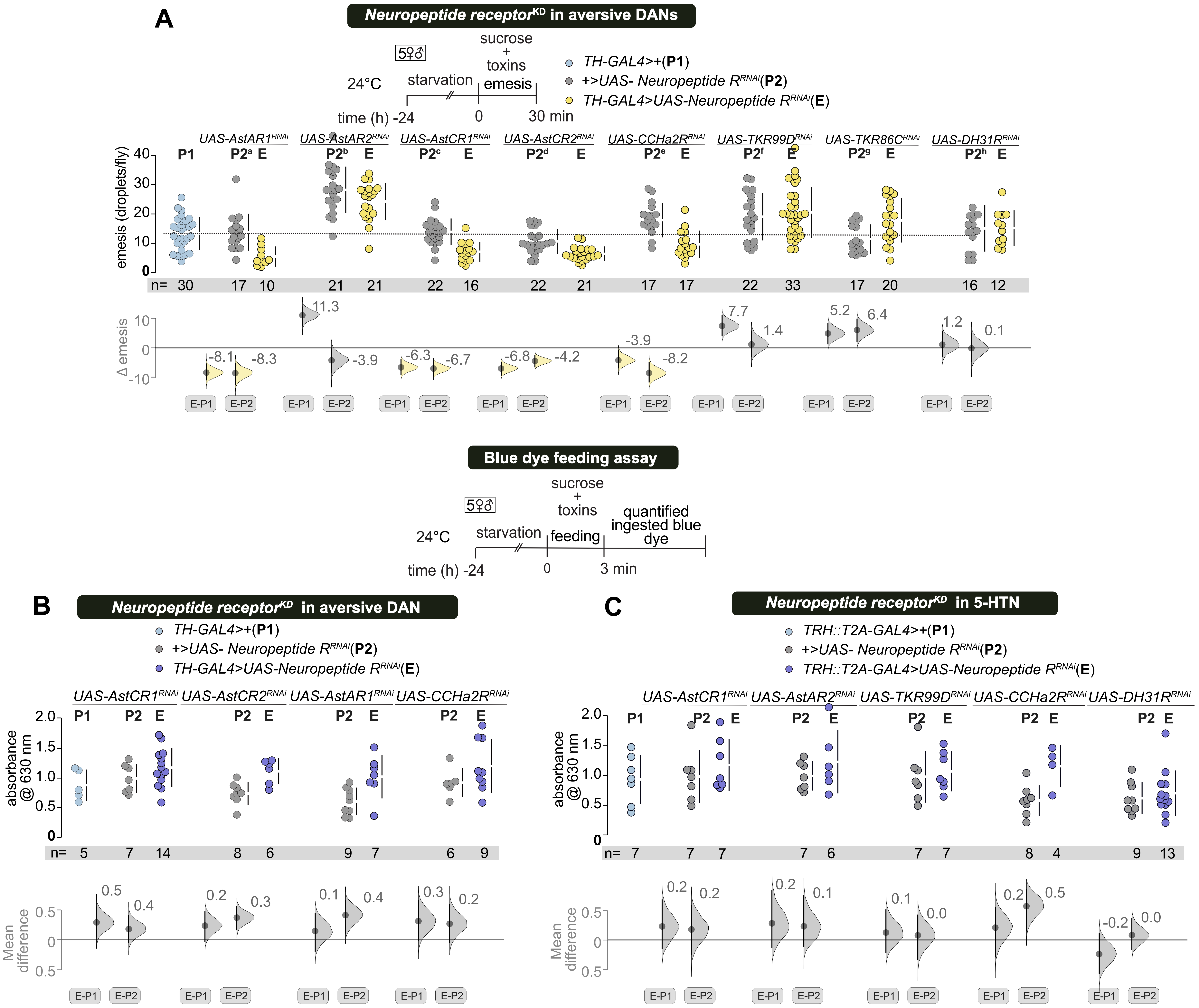
A
Neuropeptide receptor^KD in aversive DANs
24°C
starvation
sucrose + toxins
emesis
time (h) -24
0
30 min
TH-GAL4>+(P1)
+>UAS- Neuropeptide R^RNAi(P2)
TH-GAL4>UAS-Neuropeptide R^RNAi(E)
UAS-AstAR1^RNAi
UAS-AstAR2^RNAi
UAS-AstCR1^RNAi
UAS-AstCR2^RNAi
UAS-CCHa2R^RNAi
UAS-TKR99D^RNAi
UAS-TKR86C^RNAi
UAS-DH31R^RNAi
emesis (droplets/fly)
n= 30 17 10 21 21 22 16 22 21 17 17 22 33 17 20 16 12
Δ emesis
-8.1 -8.3 11.3 -3.9 -6.3 -6.7 -6.8 -4.2 -3.9 -8.2 7.7 1.4 5.2 6.4 1.2 0.1
E-P1
E-P2
Blue dye feeding assay
24°C
starvation
sucrose + toxins
feeding
quantified ingested blue dye
time (h) -24
0
3 min
B
Neuropeptide receptor^KD in aversive DAN
TH-GAL4>+(P1)
+>UAS- Neuropeptide R^RNAi(P2)
TH-GAL4>UAS-Neuropeptide R^RNAi(E)
UAS-AstCR1^RNAi
UAS-AstCR2^RNAi
UAS-AstAR1^RNAi
UAS-CCHa2R^RNAi
absorbance @ 630 nm
n= 5 7 14 8 6 9 7 6 9
Mean difference
0.5 0.4 0.2 0.3 0.1 0.4 0.3 0.2
C
Neuropeptide receptor^KD in 5-HTN
TRH::T2A-GAL4>+(P1)
+>UAS- Neuropeptide R^RNAi(P2)
TRH::T2A-GAL4>UAS-Neuropeptide R^RNAi(E)
UAS-AstCR1^RNAi
UAS-AstAR2^RNAi
UAS-TKR99D^RNAi
UAS-CCHa2R^RNAi
UAS-DH31R^RNAi
absorbance @ 630 nm
n= 7 7 7 7 6 7 7 8 4 9 13
Mean difference
0.2 0.2 0.2 0.1 0.1 0.0 0.2 0.5 -0.2 0.0

**Figure S5-1. Neuropeptide modulates aversive DANs to induce emesis**

**(A)** Knockdown of *AstAR1*, *AstCR1*, *AstCR2*, and *CCHa2R* in *TH-GAL4* labeled dopaminergic neurons (DANs) resulted in a moderate reduction in emesis, while knockdown of *AstA-R2*, *TK99D*, *TK86C*, and *DH31R* had minimal effects. **(B, C)** Knockdown of *AstAR1*, *AstCR1*, *AstCR2*, and *CCHa2R* in *TH-GAL4* and *AstAR2*, *AstCR1*, *CCHa2R* and *DH31R*, *TK99D* in *TRH::T2A-GAL4* did not impair short-term feeding measured by blue-dyed food ingestion.

### Figure -S6-1

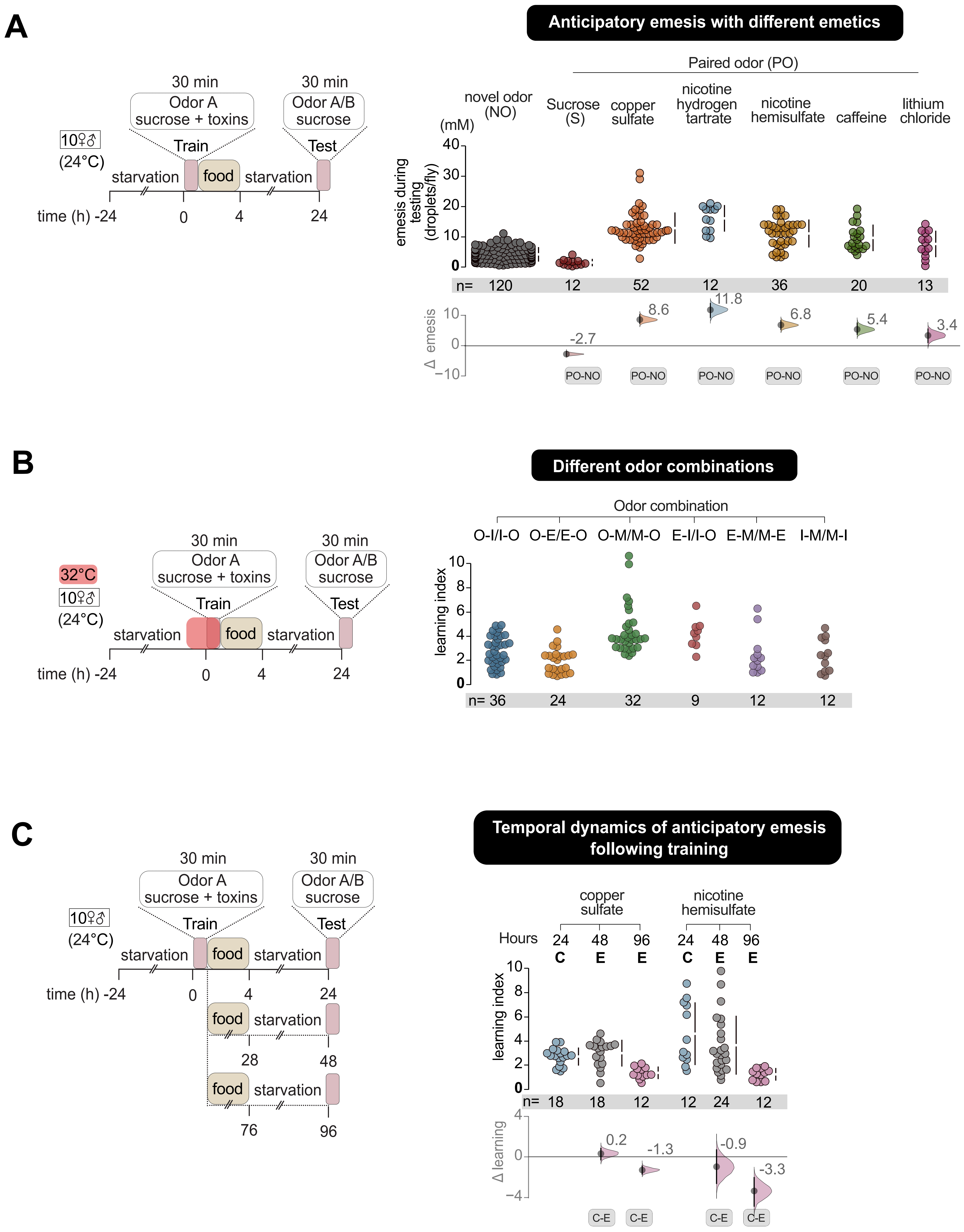
A
Anticipatory emesis with different emetics
30 min
Odor A
sucrose + toxins
30 min
Odor A/B
sucrose
Train
Test
10♀♂
(24°C)
starvation
food
starvation
time (h) -24
0
4
24
Paired odor (PO)
novel odor (NO)
Sucrose (S)
copper sulfate
nicotine hydrogen tartrate
nicotine hemisulfate
caffeine
lithium chloride
(mM)
emesis during testing (droplets/fly)
n= 120 12 52 12 36 20 13
Δ emesis
-2.7
8.6
11.8
6.8
5.4
3.4
PO-NO
B
Different odor combinations
32°C
10♀♂
(24°C)
Odor combination
O-I/I-O O-E/E-O O-M/M-O E-I/I-O E-M/M-E I-M/M-I
learning index
n= 36 24 32 9 12 12
C
Temporal dynamics of anticipatory emesis following training
copper sulfate
nicotine hemisulfate
Hours 24 48 96 24 48 96
C E E C E E
n= 18 18 12 12 24 12
Δ learning
0.2
-1.3
-0.9
-3.3
C-E
28
48
76
96

**Figure S6-1. Anticipatory emesis memory persists for 48 hours**

**(A)** Mixed sex flies exhibit anticipatory emesis in response to multiple toxins and bitter-tasting compounds. Flies trained with emetic compounds (copper sulfate, nicotine salts, caffeine, and lithium chloride) exhibited significant aversive experience indices compared to sugar fed flies. **(B)** anticipatory emesis with multiple odor pairs. To exclude odor-specific effects, flies were tested with different odor combinations. Based on these results, octan-3-ol (OCT) and methylcyclohexanol (MCH) were used as the standard odor pair in subsequent learning experiments. **(C)** Decline in anticipatory emesis over time. Anticipatory emesis memory reinforced with copper sulfate or a nicotine salt declines between 2 and 4 days post-training.

### Figure -S6-2

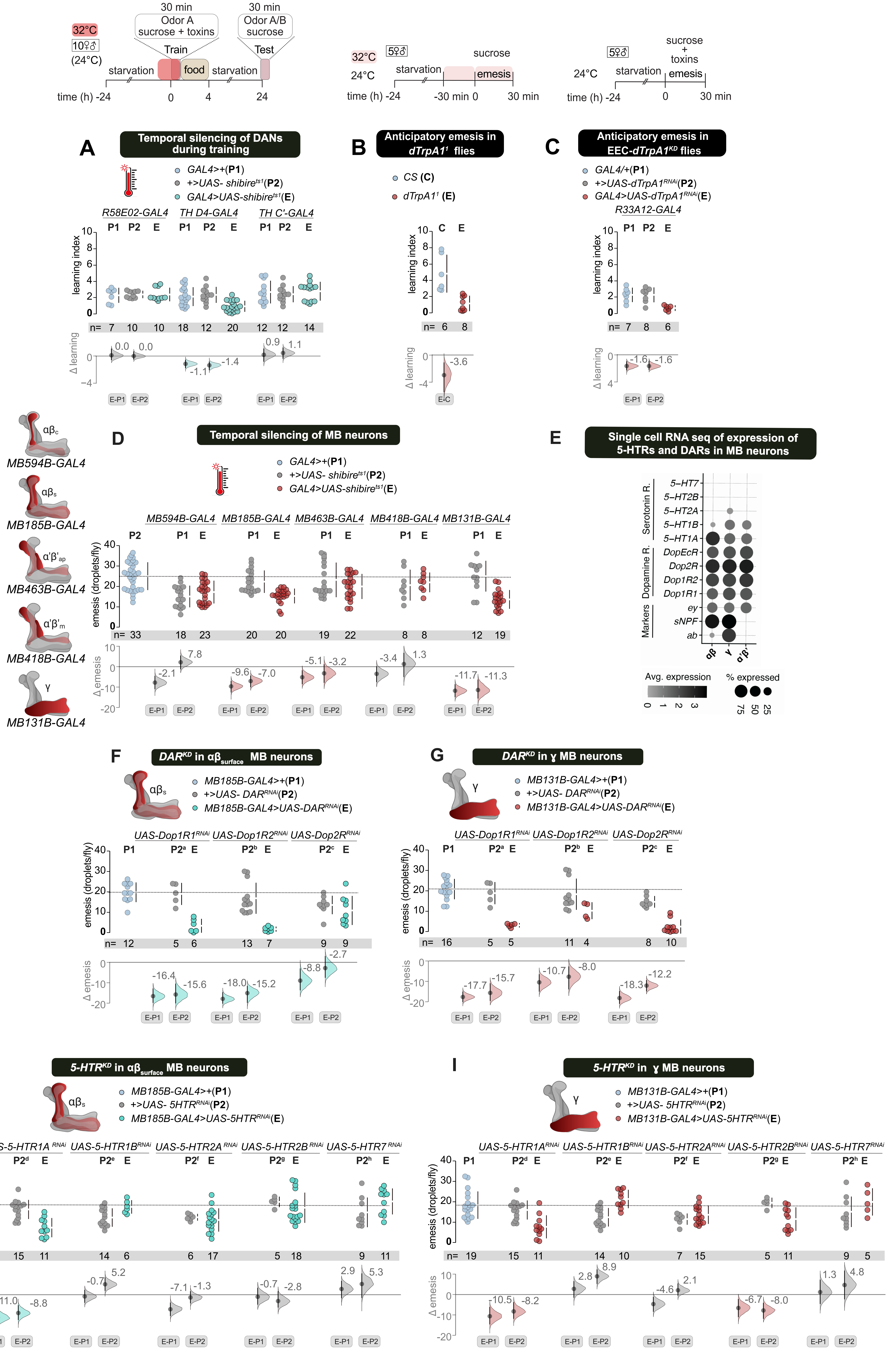

30 min
Odor A
sucrose + toxins
Train
30 min
Odor A/B
sucrose
Test
32°C
10♀♂
(24°C)
starvation
food
time (h) -24
0
4
24
5♀♂
24°C
sucrose
emesis
-30 min 0
30 min
sucrose + toxins
A
Temporal silencing of DANs during training
GAL4>+(P1)
+>UAS- shibire^ts1(P2)
GAL4>UAS-shibire^ts1(E)
R58E02-GAL4
TH D4-GAL4
TH C'-GAL4
learning index
Δ learning
B
Anticipatory emesis in dTrpA1^1 flies
CS (C)
dTrpA1^1 (E)
C
Anticipatory emesis in EEC-dTrpA1^KD flies
GAL4/+(P1)
+>UAS-dTrpA1^RNAi(P2)
GAL4>UAS-dTrpA1^RNAi(E)
R33A12-GAL4
D
Temporal silencing of MB neurons
MB594B-GAL4
MB185B-GAL4
MB463B-GAL4
MB418B-GAL4
MB131B-GAL4
emesis (droplets/fly)
Δ emesis
E
Single cell RNA seq of expression of 5-HTRs and DARs in MB neurons
Serotonin R.
Dopamine R.
Markers
5-HT7
5-HT2B
5-HT2A
5-HT1B
5-HT1A
DopEcR
Dop2R
Dop1R2
Dop1R1
ey
sNPF
ab
Avg. expression
% expressed
F
DAR^KD in αβ_surface MB neurons
MB185B-GAL4>+(P1)
+>UAS- DAR^RNAi(P2)
MB185B-GAL4>UAS-DAR^RNAi(E)
UAS-Dop1R1^RNAi
UAS-Dop1R2^RNAi
UAS-Dop2R^RNAi
G
DAR^KD in γ MB neurons
MB131B-GAL4>+(P1)
+>UAS- DAR^RNAi(P2)
MB131B-GAL4>UAS-DAR^RNAi(E)
H
5-HTR^KD in αβ_surface MB neurons
MB185B-GAL4>+(P1)
+>UAS- 5HTR^RNAi(P2)
MB185B-GAL4>UAS-5HTR^RNAi(E)
UAS-5-HTR1A^RNAi
UAS-5-HTR1B^RNAi
UAS-5-HTR2A^RNAi
UAS-5-HTR2B^RNAi
UAS-5-HTR7^RNAi
I
5-HTR^KD in γ MB neurons
MB131B-GAL4>+(P1)
+>UAS- 5HTR^RNAi(P2)
MB131B-GAL4>UAS-5HTR^RNAi(E)

**Figure S6-2: 5-HTNs and DANs tune MB neurons to regulate anticipatory emesis**

**(A)** Transient silencing using *UAS-shibire*$^{ts1}$ of DANs labeled by *TH-D4 GAL4*, which includes the PPL1, PPM2, and PPM3 clusters during training, reduces anticipatory emesis. In contrast, silencing DANs labeled by *R58E02-GAL4* and *TH-C' GAL4*, which do not include the PPL1 cluster, does not affect anticipatory emesis. **(B)** *dTrpA1* mutant (*dTrpA1*$^{1}$) flies trained with a mixture of sugar and emetics in the presence of an odor show impaired anticipatory emesis after 24 hours. **(C)** RNAi-mediated knockdown of *dTrpA1* in EECs labeled by *R33A12-GAL4* flies results in impaired anticipatory emesis. **(D)** Transient synaptic blockade of MB αβ-surface neurons labeled by *MB185B-GAL4* and γ-main neurons labeled by *MB131B-GAL4* using *UAS-shibire*$^{ts1}$ impairs emesis. **(E)** Single-cell transcriptomic data (Croset et al., 2018) show the expression of 5-HT receptors (5-HTRs) and dopamine receptors (DARs) on MB neurons. **(F, G)** RNAi-mediated knockdown of *Dop1R1* and *Dop1R2* in αβ-surface neurons (*MB185B-GAL4*) and γ-main neurons (*MB131B-GAL4*) impairs emesis. Additionally, knockdown of *Dop2R* in γ-main neurons (*MB131B-GAL4*) also impairs emesis. **(H, I)**. RNAi-mediated knockdown of *5-HTR1A* in MB αβ-surface neurons (*MB185B-GAL4*) and of *5-HTR1A* and *5-HTR2B* in MB γ-main neurons (*MB131B-GAL4*) decreases emesis, while other 5-HTRs show no effect.

#### Supplementary Table 1: Transgenic flies used in this study

| Short name | Complete genotype | Source/Reference |
|---|---|---|
| Canton-S | | Lab stock |
| $w^{1118}$ | | Gift from Shahnaz Lone |
| TrpA1 mutant | *Drosophila melanogaster:* *w[1118]; TI{w[+mW.hs]=TI}TrpA1[1]* | BDSC #26504 |
| Gr66a-GAL4 | NA | Scott et al., 2001 |
| Gr33a-RNAi | *Drosophila melanogaster:* *y[1] v[1]; P{y[+t7.7] v[+t1.8]=TRiP.HMJ30017}attP40* | BDSC #62940 |
| Gr66a-RNAi | *Drosophila melanogaster:* *y[1] v[1]; P{y[+t7.7] v[+t1.8]=TRiP.JF01225}attP2* | BDSC #31284 |
| UAS- dTrpA1 | NA | Hamada et al., 2008 |
| UAS-Shibire$^{ts1}$ | *Drosophila melanogaster:* *uas-shibire [ts1] (on X and III)* | Kitamoto, 2001 |
| UAS-Kir2.1 | *Drosophila melanogaster:* w[*];P{w[+mC]=UAS-Hsap\KCNJ2.EGFP}1 | BDSC #6596 |
| UAS- mCD8::GFP | *Drosophila melanogaster* *w[1118]; P{y[+t7.7] w[+mC]=10XUAS-IVS-mCD8::GFP}su(Hw)attP1* | BDSC #32187 |
| UAS-DenMark:: mcherry dSyt::GFP | *Drosophila melanogaster:* *w[1118]; L[1]/CyO; P{w[+mC]=UAS-DenMark}3, P{w[+mC]=UAS-syt.eGFP}3* | BDSC #33065 |
| UAS-nls-GFP | NA | Gift from Girish Ratnaparkhi |
| UAS-nSyb- GAL80 | *Drosophila melanogaster:* | BDSC #92153 |

| | | |
|---|---|---|
| | *P{w[+mC]=nSyb-GAL80.S}1, w[*]* | |
| Double balancer 2nd and 3rd chromosome | *Drosophila melanogaster:*<br>*w; sp/Cyo; Dr/TM3sb* | VDRC #v51427 |
| Double balancer 1st and 3rd chromosome | *Drosophila melanogaster:*<br>*FM7a;;MKRS/TB* | Gift from Girish Ratnaparkhi |
| R58E02-GAL4 | *Drosophila melanogaster:*<br>w[1118]; P{y[+t7.7] w[+mC]=GMR58E02-GAL4}attP2 | BDSC #41347 |
| TH-GAL4 | *Drosophila melanogaster:*<br>w[*]; P{w[+m*]=ple-GAL4.F}2/CyO | BDSC #95269 |
| TH-C'-GAL4 | *Drosophila melanogaster:*<br>*w[*]; P{w[+mC]=ple-GAL4.TH-C'}3/TM6C, Sb[1]* | BDSC #93703 |
| TH-D'-GAL4 | *Drosophila melanogaster:*<br>*w[*]; P{w[+mC]=ple-GAL4.TH-D'}2/CyO* | BDSC #93704 |
| TH-D4-GAL4 | NA | Liu et al., 2012 |
| MB296B-GAL4 | *Drosophila melanogaster:*<br>*w[1118]; P{y[+t7.7] w[+mC]=R15B01-p65.AD}attP40; P{y[+t7.7] w[+mC]=R26F01-GAL4.DBD}attP2* | BDSC #68308 |
| MB058B-GAL4 | *Drosophila melanogaster:*<br>*w[1118]; P{y[+t7.7] w[+mC]=R82C10-p65.AD}attP40; P{y[+t7.7] w[+mC]=R50B03-GAL4.DBD}attP2* | BDSC #68278 |
| MB320C-GAL4 | *Drosophila melanogaster:*<br>*w[1118]; P{y[+t7.7] w[+mC]=R22B12-GAL4.DBD}attP2 PBac{y[+mDint2] w[+mC]=ple-p65.AD}VK00027* | BDSC #68253 |

| | | |
|---|---|---|
| UAS-Dop1R1 RNAi | *Drosophila melanogaster:* *y[1] sc[*] v[1] sev[21]; P{y[+t7.7] v[+t1.8]=TRiP.HMC02344}attP2/TM, Sb[1]* | BDSC #55239 |
| UAS-Dop1R2 RNAi | NA | Gift from Sheeba Vasu |
| TRH-Gal4 | NA | Alekseyenko et al., 2010 |
| TRH::T2A-Gal4 | *Drosophila melanogaster:* *w[*]; TI{2A-GAL4}Trhn[2A-GAL4]* | BDSC #84694 |
| TRH-K2-GAL4 | *Drosophila melanogaster:* *w[1118]; P{w[+mC]=Trhn-GAL4.long}2* | BDSC #38388 |
| TRH-K3-GAL4 | *Drosophila melanogaster:* *w[1118]; P{w[+mC]=Trhn-GAL4.long}3* | BDSC #38389 |
| VT064246-GAL4 | NA | VDRC #204311 |
| UAS-TRH RNAi | *Drosophila melanogaster:* *y[1] v[1]; P{y[+t7.7] v[+t1.8]=TRiP.JF01863}attP2/TM3, Sb[1]* | BDSC #25842 |
| UAS-SerT RNAi | NA | BDSC #62985 |
| UAS-TH RNAi | NA | NA |
| UAS-5-HTR1A RNAi | *Drosophila melanogaster:* *y[1] sc[*] v[1] sev[21]; P{y[+t7.7] v[+t1.8]=TRiP.HMS00823}attP2* | BDSC #33885 |
| UAS-5-HTR1B RNAi | NA | VDRC #v35791 |
| UAS-5HTR2A RNAi | NA | Gift from Sheeba Vasu VDRC #v102105 |
| UAS 5-HTR2B RNAi | *Drosophila melanogaster:* *y[1] v[1]; P{y[+t7.7] v[+t1.8]=TRiP.HMJ22882}attP40* | BDSC #60488 |

| | | |
|---|---|---|
| UAS 5-HTR7 RNAi | NA | Gift from Sheeba Vasu |
| MB131B-GAL4 | *Drosophila melanogaster:*<br><br>*w[1118]; P{y[+t7.7] w[+mC]=R13F02-p65.AD}attP40/CyO; P{y[+t7.7] w[+mC]=R89B01-GAL4.DBD}attP2* | BDSC #68265 |
| MB594B-GAL4 | *Drosophila melanogaster:*<br><br>*w[1118]; P{y[+t7.7] w[+mC]=R13F02-p65.AD}attP40; P{y[+t7.7] w[+mC]=R58F02-GAL4.DBD}attP2* | BDSC #68255 |
| MB185B-GAL4 | *Drosophila melanogaster:*<br><br>*w[1118]; P{y[+t7.7] w[+mC]=R52H09-p65.AD}attP40; P{y[+t7.7] w[+mC]=R18F09-GAL4.DBD}attP2* | BDSC #68267 |
| MB418-GAL4 | *Drosophila melanogaster:*<br><br>*w[1118]; P{y[+t7.7] w[+mC]=R26E07-p65.AD}attP40/CyO; P{y[+t7.7] w[+mC]=R30F02-GAL4.DBD}attP2* | BDSC #68322 |
| UAS-TrpA1 RNAi | *Drosophila melanogaster:*<br><br>*y[1] v[1]; P{y[+t7.7] v[+t1.8]=TRiP.JF01360}attP2* | BDSC #31384 |
| UAS-TK RNAi | *Drosophila melanogaster:*<br><br>*y[1] v[1]; P{y[+t7.7] v[+t1.8]=TRiP.JF01818}attP2* | BDSC #25800 |
| UAS-AstC RNAi | *Drosophila melanogaster:*<br><br>*y[1] v[1]; P{y[+t7.7] v[+t1.8]=TRiP.JF01907}attP2* | BDSC #25868 |
| UAS-AstA RNAi | *Drosophila melanogaster:*<br><br>*y[1] v[1]; P{y[+t7.7] v[+t1.8]=TRiP.JF01905}attP2* | BDSC #25866 |
| UAS-DH31 RNAi | *Drosophila melanogaster:* | BDSC #41957 |

| | | |
|---|---|---|
| | *y[1] sc[*] v[1] sev[21]; P{y[+t7.7] v[+t1.8]=TRiP.HMS02354}attP2/TM3 , Sb[1]* | |
| UAS-CCHa1 RNAi | *Drosophila melanogaster:*<br><br>*y[1] sc[*] v[1] sev[21]; P{y[+t7.7] v[+t1.8]=TRiP.HMC04879}attP40* | BDSC #57562 |
| UAS-CCHa2 RNAi | *Drosophila melanogaster:*<br><br>*y[1] sc[*] v[1] sev[21]; P{y[+t7.7] v[+t1.8]=TRiP.HMC04565}attP40* | BDSC #57183 |
| UAS-NPF RNAi | *Drosophila melanogaster:*<br><br>*y[1] v[1]; P{y[+t7.7] v[+t1.8]=TRiP.JF02555}attP2* | BDSC #27237 |
| UAS-TkR86C RNAi | *Drosophila melanogaster:*<br><br>*y[1] v[1]; P{y[+t7.7] v[+t1.8]=TRiP.JF02160}attP2* | BDSC #31884 |
| UAS-TkR99D RNAi | *Drosophila melanogaster:*<br><br>*y[1] sc[*] v[1] sev[21]; P{y[+t7.7] v[+t1.8]=TRiP.HMC03749}attP40* | BDSC #55732 |
| UAS-AstCR1 RNAi | *Drosophila melanogaster:*<br><br>*y[1] v[1]; P{y[+t7.7] v[+t1.8]=TRiP.JF02656}attP2/TM3, Sb[1]* | BDSC #27506 |
| UAS-AstCR2 RNAi | *Drosophila melanogaster:*<br><br>*y[1] v[1]; P{y[+t7.7] v[+t1.8]=TRiP.JF01960}attP2* | BDSC #25940 |
| UAS-AstAR1 RNAi | *Drosophila melanogaster:*<br><br>*y[1] v[1]; P{y[+t7.7] v[+t1.8]=TRiP.JF02578}attP2* | BDSC #27280 |
| UAS-AstAR2 RNAi | *Drosophila melanogaster:*<br><br>*y[1] sc[*] v[1] sev[21]; P{y[+t7.7] v[+t1.8]=TRiP.HMS05683}attP40* | BDSC #67864 |
| UAS-CCHa2R RNAi | *Drosophila melanogaster:*<br><br>*y[1] v[1]; P{y[+t7.7] v[+t1.8]=TRiP.JF01876}attP2* | BDSC #25855 |

| | | |
|---|---|---|
| NP1-Gal4 | NA | Jiang and Edgar, 2009 |
| R33A12-Gal4 | *Drosophila melanogaster:*<br><br>*w[1118]; P{y[+t7.7] w[+mC]=GMR33A12-GAL4}attP2* | BDSC #49739 |
| R34E04-Gal4 | *Drosophila melanogaster:*<br><br>*w[1118]; P{y[+t7.7] w[+mC]=GMR34E04-GAL4}attP2* | BDSC #49788 |
| R46B05-Gal4 | *Drosophila melanogaster:*<br><br>*w[1118]; P{y[+t7.7] w[+mC]=GMR46B05-GAL4}attP2* | BDSC #50253 |
| UAS-Duox RNAi | *Drosophila melanogaster:*<br><br>*y[1] sc[*] v[1] sev[21]; P{y[+t7.7] v[+t1.8]=TRiP.HMS00692}attP2* | BDSC #32903 |
| UAS-Nox RNAi | *Drosophila melanogaster:*<br><br>*y[1] sc[*] v[1] sev[21]; P{y[+t7.7] v[+t1.8]=TRiP.HMS00429}attP2* | BDSC #32433 |
| R13G06-Gal4 | *Drosophila melanogaster:*<br><br>*w[1118]; P{y[+t7.7] w[+mC]=GMR13G06-GAL4}attP2* | BDSC #48584 |
| UAS-AstB RNAi | *y[1] sc[*] v[1] sev[21]; P{y[+t7.7] v[+t1.8]=TRiP.HMS02244}attP2* | BDSC #41680 |
| UAS-amon RNAi | *Drosophila melanogaster:*<br><br>*y[1] v[1]; P{y[+t7.7] v[+t1.8]=TRiP.HM05071}attP2* | BDSC #28583 |
| MB630B-GAL4 | *Drosophila melanogaster:*<br><br>*w[1118]; P{y[+t7.7] w[+mC]=VT026773-p65.AD}attP40; P{y[+t7.7] w[+mC]=R72B05-GAL4.DBD}attP2* | BDSC #68334 |
| UAS-DH31R RNAi | *Drosophila melanogaster:*<br><br>*y[1] v[1]; P{y[+t7.7] v[+t1.8]=TRiP.JF01945}attP2/TM3, Sb[1]* | BDSC #25925 |

| | | |
|---|---|---|
| UAS-DOP2R RNAi | *Drosophila melanogaster:*<br>*w[*]; P{w[+mC]=UAS-Dop2R.RNAi.D}3* | BDSC #78804 |
| MB463B-GAL4 | *Drosophila melanogaster:*<br>*w[1118]; P{y[+t7.7] w[+mC]=R35B12-p65.AD}attP40; P{y[+t7.7] w[+mC]=R34A03-GAL4.DBD}attP2/ TM6B, Tb[1]* | BDSC #68370 |
| Piezo mutant | *Drosophila melanogaster:*<br>*w[*];PBac{w[+mC]=RB5.WH5}Piezo[KO]* | BDSC #58770 |
| Bitter-SEL-Gal4 | *Drosophila melanogaster:*<br>*w[1118]; P{y[+t7.7] w[+mC]=R24F06-p65.AD}attP40/CyO, P{2xTb[1]-RFP}CyO; P{y[+t7.7] w[+mC]=R45E06-GAL4.DBD}attP2/ TM6B, Tb[1]* | BDSC #75837 |
| UAS-Lola | *Drosophila melanogaster:*<br>*UAS-Flp/SM6a; Lola>mcherry.STOP>Luc/MKRS* | Gift from Nitin Chouhan<br>Guo et al., 2017 |

#### Supplementary Table 2: Key resources table

| Resource/Reagent | Source | Identifier |
|---|---|---|
| **Chemicals** | | |
| Agar Bacteriological grade (Cas no. 9002-18-0) | Himedia | GRM026P |
| Absolute ethanol (Cas no. 64-17-5) | Merck | 107017 |
| Malt extract (Cas no. 8002-48-0) | Himedia | RM004 |
| Propionic acid (Cas no. 79-09-4) | Loba chemie pvt.ltd | Article No. 00265 |
| Methyl paraben (Cas no. 99-76-3) | Himedia | GRM1899 |
| Sucrose (Cas no. 57-50-1) | ANJ Biomedicals | Catalog no. 100314 |
| 3-Octanol (Cas no. 589-98-0) | Sigma-Aldrich | 218405 |
| 4- Methyl cyclohexanol (Cas no 589-91-3) | Sigma-Aldrich | 153095 |
| Low melting agarose (Cas no 9012-36-6 ) | MP Biomedicals | LLC catalogue no. AGAL0050 |
| Copper (II) sulphate pentahydrate (Cas no. 7758-99-8) | Himedia | SKU: PCT0104 |
| Caffeine (Cas no. 58-08-2) | Sigma-Aldrich | C7731 |
| Coumarin (Cas no. 91-64-5) | Sigma-Aldrich | C4261 |

| | | |
|---|---|---|
| Denatonium Benzoate (Cas no. 3734-33-6) | Tokyo Chemical Industry | Pdt no. D2124 |
| Nicotine hemisulfate salt (Cas no. 65-30-5) | Sigma Life Science | N1019 |
| Lithium Chloride (Cas no. 7447-41-8) | Sigma-Aldrich | L9650 |
| Nicotine hydrogen tartrate salt (Cas no. 65-31-6) | Sigma Life Science | SML1236 |
| D-luciferin potassium salt (Cas no. 115144-35-9) | Sigma-Aldrich | 50227 |
| Phosphate Buffer Salt tablets | MP Biomedicals | Catalog no. 2810305 |
| Paraformaldehyde (Cas no. 30525-89-4) | Sigma-Aldrich | 158127 |
| Triton X-100 (Cas no. 9002-93-1) | MP biomedical | SKU:02194854-CF |
| Poly L lysine (Cas no. 25988-63-0) | Sigma-Aldrich | P1399 |
| Dithiothreitol (Cas no. 3483-12-3) | Sigma-Aldrich | D9779 |
| Dimethyl sulfoxide (Cas no. 67-68-5) | Sigma-Aldrich | D8418 |
| Dihydroethidium (Cas no. 104821-25-2) | Sigma-Aldrich | D7008 |
| Brilliant blue FCF food grade (Cas no. 3844-45-9) | Vidhi Manufacturers 2490 | |

| | | |
|---|---|---|
| Glycerol (Cas no.56-81-5) | Qualigens, Thermo Fisher Scientific | |
| Table sugar | Madhur brand, refined sugar | |
| Goat serum | Sigma Aldrich | |
| Brewer’s yeast | Prime brand, instant dry yeast | |
| **Antibodies** | | |
| Serotonin Antibody (YC5/45) [Alexa Fluor® 594] | Novus Biologicals | NB100-65037AF594 |
| **Software and resources** | | |
| Prism | Graphpad | https://www.graphpad.com/ |
| BioRender | | https://www.biorender.com/ |
| Shotcut | | https://www.shotcut.org/ |
| Fiji/ImageJ | | https://imagej.net/software/fiji/ |
| Affinity | | https://affinity.serif.com/en-us/designer/ |
| Estimation stats | | https://www.estimationstats.com/ |
| Source for neurotransmitter annotations | | https://github.com/funkelab/drosophila_neurotransmitters |
| Neuron reconstruction for visualization and analysis | | https://github.com/seung-lab/cloud-volume |
| Calculating the network influence score based on the connectome calculator | | https://github.com/natverse/influencer |

#### Additional supplementary files

https://drive.google.com/drive/folders/17doUNU4XDD24UhFGGqp7yKYGcui58kHR?usp=sharing

**Supplementary Table 3: Root IDs**
**Supplementary Video 1: Latency, emesis and defecation**
**Supplementary Video 2: Crop contractions**
**Supplementary Video 3: Anticipatory emesis**
